# Integrated Blood–Brain Transcriptomics Reveals CD79A and GRIA2 as Drug Repurposing Targets in Multiple Sclerosis

**DOI:** 10.1101/2025.09.02.673787

**Authors:** Mohamed A. Mahmoud, Mohammed M. Alshehri

## Abstract

Multiple sclerosis (MS) has concomitant immune and neurodegenerative mechanisms that are not tractable to single□mechanism treatments. We speculated that simultaneous transcriptomic–network analysis of peripheral blood mononuclear cells (PBMCs) and MS brain lesions would identify convergent, druggable hubs susceptible to combination repurposing approaches. Microarray data of PBMCs (GSE21942; 14 MS, 15 controls) and brain lesions (GSE38010; 5 MS, 2 controls) were analyzed using differential□expression thresholds (adjusted P < 0.05; |log□FC| ≥ 1.5). Protein–protein□interaction networks were constructed with Search Tool for the Retrieval of Interacting Genes and subnetworks recognized by molecular complex detection. PBMC co□expression□modules were recognized by Weighted Gene Co-expression Network Analysis. Functional enrichments of cluster genes were undertaken by gene ontology and gene set enrichment analysis. CytoHubba identified candidate genes first, then hubs were identified by Least Absolute Shrinkage and Selection Operator (LASSO) regression (PBMCs) and top three brain lesion CytoHubba genes were considered hub genes. We screened a ligand library against four hubs CD79A, GRIN2A, NRXN1, and GRIA2, and assessed binding stability by 200 ns molecular dynamics and MM/PBSA. We found 142 PBMC differentially expressed genes (DEGs) and 1493 lesion DEGs, which were mapped to four peripheral modules B□cell□receptor signalling, erythrocyte metabolism, spliceosome stress and dampened innate sensing, and four central modules glutamatergic synapse, axon ensheathment/sodium□channel stress, netrin□1 signalling and nucleocytoplasmic□transport. LASSO identified FCRL1, CD22, and CD79A, while GRIN2A, GRIA2, and NRXN1 were the chosen brain hubs based on CytoHubba Maximal Clique Centrality. Molecular docking and dynamics simulation identified icotinib and niraparib as dual□target ligands of CD79A and GRIA2. Our end□to□end discovery pipeline defines CD79A and GRIA2 as dual□compartment targets. Repurposing compounds like icotinib and niraparib to co□modulate B□cell activation and excitotoxic synaptic injury represents an actionable strategy for multipronged treatments of MS.

## 1. Introduction

Multiple sclerosis (MS) is a chronic inflammatory and neurodegenerative disorder of the central nervous system, affecting approximately 2.8 million individuals worldwide and ranked among the leading causes of non□traumatic neurological disability in young adults [1]. Clinically, MS typically follows a relapsing-remitting course that transitions in many patients to a progressive phase characterized by accumulating disability and neurodegeneration despite effective suppression of focal inflammation [2, 3]. Pathologically, MS lesions display a complex interplay of peripheral immune infiltration, blood-brain barrier (BBB) breakdown, focal demyelination and secondary axonal transection, reflecting both “outside-in” autoimmune mechanisms where autoreactive T and B lymphocytes breach the BBB and target myelin and “inside-out” processes, in which primary oligodendrocyte or neuronal injury exposes CNS antigens to the periphery [4, 5].

In recent years, B cells have emerged as pivotal mediators in MS pathogenesis. Early trials of anti-CD20 monoclonal antibodies such as rituximab demonstrated a more than 50 % reduction in clinical relapse rates in relapsing-remitting MS patients [6], and subsequent studies of the Bruton’s tyrosine kinase (BTK) inhibitor tolebrutinib confirmed that B-cell receptor□axis modulation can further attenuate lesion formation in relapsing disease [7]. However, progression independent of relapse activity (PIRA) now drives most long□term disability in MS, highlighting the urgent need for treatments that target both immune and neurodegenerative processes [3].

Although transcriptomic studies have shed light on MS mechanisms, they rarely integrate peripheral and central data. By combining protein–protein interaction (PPI) networks and co□expression analysis, we can pinpoint core molecular modules across compartments [8, 9], and *in silico* drug-repurposing through virtual screening and molecular-dynamics allows us to test existing compounds against these targets quickly [10].

In this study, we hypothesized that an integrated transcriptomic–network analysis of peripheral blood mononuclear cells (PBMCs) and MS brain lesions would reveal convergent, druggable hubs amenable to combinational therapy. We analyzed microarray data from PBMCs (GSE21942; 14 MS, 15 controls) and brain lesions (GSE38010; 5 MS, 2 controls) under uniform preprocessing and differential□expression criteria (adjusted P□<□0.05; |log□fold□change| ≥ 1.5). PPI networks were constructed in the Search Tool for the Retrieval of Interacting Genes (STRING) database and subnetworks defined by molecular complex detection (MCODE), while co-expression modules were delineated via Weighted Gene Co-expression Network Analysis (WGCNA). Gene ontology and gene set enrichment analysis (GSEA) annotated functional pathways. Candidate hubs were prioritized through CytoHubba and machine□learning model LASSO regression. Finally, we performed virtual screening of a ligand library against two top hubs (CD79A in PBMCs, GRIA2 in lesions) and validated binding stability through 200-ns molecular□dynamics simulations and free binding energy calculation using MM/PBSA.

## 2. Materials and Methods

### 2.1. Data sources, processing, and identification of differentially expressed genes

Gene expression data sets GSE21942 platform GPL570 (PBMCs: 14 MS vs. 15 controls) and GSE38010 platform GPL570 (brain tissue: 5 MS lesions vs. 2 controls) were obtained from the NCBI Gene Expression Omnibus through the GEOquery R package [11]. Expression matrices were downloaded through Biobase and log□-transformed. Quantile normalization was carried out with limma normalizeBetweenArrays for making inter-sample comparable [12, 13]. For quality control, boxplots were used to determine distribution uniformity, principal component analysis plots to determine group discrimination, and Pearson correlation heatmaps to determine intra- and inter-group correlations. The grouping variables were specified and Surrogate Variable Analysis (SVA) using the svaseq function from the sva R package was employed to account for potential batch effects [14]. Eight significant surrogate variables were identified for GSE21942 and two for GSE38010 and were subsequently incorporated into the linear modeling design matrix. Linear modeling was carried out through the limma package, precision weights having been estimated through arrayWeights, and empirical Bayes moderation was applied [15]. DEGs were considered genes having an adjusted p-value < 0.05 (Benjamini-Hochberg correction) and |log□fold change| ≥ 1.5. Volcano plots and heatmaps were generated using ggplot2, ggrepel, and pheatmap, and the top 10 up- and downregulated DEGs were tagged for interpretation purposes [16].

### 2.2. Protein-protein interactions, identification of candidate genes, and functional module analysis

The predicted protein-protein interactions of the significant DEGs (adjusted p-value < 0.05 (Benjamini-Hochberg correction) and |log fold change| ≥ 1.5) of both GSE21942 and GSE38010 were retrieved from the STRING database (https://string-db.org/) [17]. The official gene symbols were inserted under the “Multiple Proteins” with organism set to “Homo Sapiens”. The full STRING networks were obtained using a medium confidence. Next, the attained PPI networks were exported to Cytoscape software (version 3.10.1, Boston, MA, USA) [18]. The top 10 genes in the attained networks were identified using CytoHubba according to the Maximal Clique Centrality option. The attained CytoHubba genes of the networks were considered as the candidate genes for subsequent bioinformatics analysis except the top three genes of brain lesion were considered as hub genes. The MCODE algorithm of Cytoscape software (version 3.10.1, Boston, MA, USA) was used to uncover the subnetworks with settings set as follows (degree cut-off = 2, node score cut-off = 0.2, k-core = 2, and maximum depth = 100) [19]. To visualize the protein–protein interaction networks and highlight functional modules and candidate genes, edge and node attribute tables were exported from Cytoscape following STRING-based network construction, CytoHubba hub gene ranking, and MCODE clustering. These tables were imported into R and processed using the igraph, tidygraph, and ggraph packages [20, 21]. The igraph package was used to build undirected network graphs using graph_from_data_frame with annotated node and edge information. For publication-quality visualization, the networks were converted into tidy graph structures with tbl_graph from the tidygraph package and plotted using ggraph, applying Fruchterman–Reingold force-directed layouts. Node colours were mapped to log fold change. Edge transparency and width were scaled to STRING confidence scores.

### 2.3. Gene enrichment analysis and gene set enrichment analysis

In the PBMCs data set (GSE21942), DEGs of every cluster identified with MCODE in the PPI network were retrieved and converted to Entrez IDs with the clusterProfiler package’s bitr function using the org.Hs.eg.db annotation database [22]. GO enrichment of Biological Process (BP), Cellular Component (CC), and Molecular Function (MF) was conducted via the enrichGO function and Reactome pathway enrichment was conducted via enrichPathway function from ReactomePA package with significance thresholds set at p < 0.05 and q < 0.2 using Benjamini-Hochberg correction [23]. For the brain data set (GSE38010), each of the four resulting MCODE clusters were also analyzed by the identical GO and Reactome pathway enrichment pipeline. We further conducted GSEA for all GSE38010 DEGs ranked by log□fold change using the gseGO and gsePathway functions for GO and Reactome enrichment and GSEA function for MSigDB-derived WikiPathways using the msigdbr package with parameters including 1000 permutations, gene set sizes between 10 and 500, and a p-value cutoff of 0.05 with Benjamini-Hochberg adjustment [24]. Enrichment plots were plotted to provide GSEA results for both upregulated and downregulated gene sets in the three databases.

### 2.4. Weighted gene co-expression network analysis

To show co-expressed clusters of genes and analyze their association with disease status, we carried out WGCNA with the R package WGCNA [8, 25]. To collapse redundant probes, we calculated the variance of each probe across all samples and, after ranking in descending order, retained for each gene symbol only the probe with the highest variance; all other duplicates were removed. The resulting deduplicated matrix was normalized across arrays by quantile normalization to correct for technical differences between samples. We then computed per-gene variance in the normalized data and excluded the lowest 20 percent of genes by this metric, thereby focusing the analysis on the most informative genes. Quality control was performed via goodSamplesGenes, which flagged and removed any samples or genes exhibiting excessive missing values or zero variance. We initiated the calculation of pair-wise Pearson correlations between all the genes from the samples to generate a similarity matrix. To highlight significant correlations and eliminate noise, we used a soft-thresholding power of 4, selected using the pickSoftThreshold function, based on the criterion of approximate scale-free topology. The resultant adjacency matrix was converted to a Topological Overlap Matrix (TOM), which contains direct and indirect interaction between genes. Genes were hierarchically clustered according to TOM-based dissimilarity, and co-expressed modules of genes were determined by the dynamic tree cut with parameters (deepSplit = 2, minModuleSize = 30, and mergeCutHeight = 0.25) using the blockwiseModules function, resulting in multiple gene modules, each labelled with a unique colour [8, 26]. Module eigengenes (first principal component of gene expression within a module) were computed and correlated with disease status to determine MS-related modules. We produced a heatmap to observe module–disease associations and chose modules that were strongly correlated with the trait for closer examination. For every one of these modules, we chose the top 10 hub genes based on their intramodular connectivity. The top 10 most significant hub genes from every module were presented by employing the igraph package in R. For each significant WGCNA module, we pulled out its member genes, converted their symbols to Entrez IDs, and ran a GO Biological Process over-representation test (Benjamini-Hochberg-adjusted p□<□0.05). We then selected the top ten enriched terms.

### 2.5. LASSO regression

To find the hub genes, we employed predictive modeling through machine learning and feature selection on the top 10 candidate genes selected by the CytoHubba plugin in Cytoscape for the data set of PBMCs (GSE21942). We used LASSO regression with the glmnet R package to PBMCs data with stratified five-fold cross-validation to determine the best penalty parameter lambda that would give the best area under the curve (AUC) [27–29]. From that, we picked the simplest model whose performance was within one standard error of the best, and kept only the genes with non-zero coefficients in that model. We then combined all the held-out predictions to plot a receiver operating characteristic (ROC) curve for the selected model using the pROC package [30]. Finally, to obtain an unbiased performance estimate, we used nested cross-validation (five outer folds for model evaluation, and within each outer training set, a five-fold inner cross-validation) and applied it to the held-out outer fold and we calculated the ROC AUC.

### 2.6. Preparation of ligands and proteins

The X-ray structures of CD79A (PDB ID: 7XQ8), GRIN2A (PDB ID: 5KCJ), NRXN1 (PDB ID: 3B3Q), and GRIA2 (PDB ID: 5YBG) were downloaded from the Research Collaboratory for Structural Bioinformatics Protein Data Bank (https://www.rcsb.org/).The proteins were prepared using University of California San Francisco Chimera (https://www.cgl.ucsf.edu/chimera/) [31]. Water and heteroatoms were removed, and gaps in residues, side chains, and loops were fixed. All hydrogens, such as non-polar hydrogens and Gasteiger charges, were added to protein structures. Partial charges were distributed on missing amino acid residue atoms to ensure system charge consistency. Moreover, a library of nearly 2,500 ligands was obtained from Selleckchem (https://www.selleckchem.com/) and the ligands were subjected to Lipinski’s rule of five [32], Veber’s rule [33], Egan’s rule, and Abbot’s bioavailability score. This was achieved via the web tool SwissADME (http://www.swissadme.ch/index.php) [34]. Only compounds with high BBB permeability, good bioavailability, and not violating any of the abovementioned rules were chosen to undergo virtual screening with the targets. The AI Drug Lab (https://ai-druglab.smu.edu/admet) was utilized for the prediction of the ADMET profiles of the chosen compounds [35].

### 2.7. Virtual screening and molecular docking analysis

The ligand binding site was predicted using PASSer (https://passer.smu.edu/) [36]. A receptor grid generation tool was utilized to produce the grid on the binding site of the proteins. The receptor grid shows the area where a ligand and a protein interact. This was done by choosing the top-ranked site predicted by PASSer, which reveals the binding area. Molecular docking was performed using PyRx (https://pyrx.sourceforge.io/) for multiple ligands [37]. The prepared compounds and proteins were docked to predict compounds with the least binding energy (kcal/mol). The most promising compounds were chosen to perform a molecular docking refinement as well as subsequent molecular dynamics simulations. BIOVIA was used to generate 2D interaction plots ( https://www.3ds.com/products/biovia/discovery-studio).

### 2.8. Molecular dynamics simulation analysis

Protein-ligand complex was simulated by an all-atom molecular dynamics simulation. Preparation of the simulation was done via the CHARMM-GUI Server using the solution builder protocol. The protein was protonated at physiological pH level 7.4. Ligand parameterization was achieved using software like CGenFF, while protein parameterization used the CHARMM36 force field. The system was solvated with the TIP3P water model in a periodic cubic box, which was extended by 10 Å outside the protein. 0.15 M counter ions in 73 sodium and 85 chloride ions form were added to neutralize the system. In electrostatic and van der Waals interaction simulations, we employed the Verlet cutoff method. Bond lengths within the system were kept constant using the LINCS algorithm. The particle mesh Ewald method was used to obtain precise long-range electrostatic potentials. The steepest descent algorithm was used to energy-minimize the system in order to remove unwanted contacts and decrease potential energy below 1000 kJ mol□^1^ nm□^1^. After energy minimization, the system was then subjected to two-stage equilibration. The first one was carried out in the NVT ensemble at constant temperature under a thermostat, whereas the second one used the NPT ensemble to hold both pressure and temperature in control. The temperature was kept at 310.05 K during the simulation, with the pressure kept at 1 bar to enable the system to exchange energy as well as particles. This enabled it to become thermodynamically equal with homogeneous thermodynamic properties. Radius of gyration (Rg), hydrogen bonds, solvent accessible surface area (SASA), root mean square deviation (RMSD), and root mean square fluctuation (RMSF) were some parameters that were calculated from the trajectory obtained using the GROMACS software (Version 2024.4) (https://www.gromacs.org/) [38]. XMGrace (https://ambermd.org/tutorials/Plotting.php) was utilized to plot each of the plots . MD simulation was carried out for 200 ns.

### 2.9. Binding free energy calculation

To acquire more profound understanding of the energetic driving forces behind ligand binding, we performed MM/PBSA calculations on a carefully screened set of MD snapshots [39, 40]. The free energy of binding (ΔGbind) was decomposed into discrete terms of energy: van der Waals interactions (ΔEvdW), electrostatic contributions (ΔEelec), polar solvation effects (ΔGpol), and nonpolar solvation factors (ΔGnpolar). In addition, per-residue decomposition analysis was performed to identify the important residues participating in ligand binding and stabilization. From this fine-scale analysis, the important role of single amino acid interactions in affinity was highlighted, shedding light on the complex chemical environment stabilizing the protein-ligand complex.

## 3. Results

### 3.1. PBMCs

#### 3.1.1. Identification of PBMCs DEGs

PBMCs gene expression profiles from dataset GSE21942 was preprocessed and analyzed to identify the DEGs between healthy controls and MS patients. After log_2_ transformation and quantile normalization, the boxplot visualization showed uniformly distributed expression values across all samples, with no apparent outliers, validating the effectiveness of the preprocessing pipeline (**Figure 1A**). Principal component analysis clearly demonstrated segregation of control and MS groups and thus different transcriptional signatures (**Figure 1B**). This was further supported by hierarchical clustering of Pearson correlation coefficients, which showed strong intra-group similarity and robust inter-group separation with the lowest correlation value nearing 0.975 (**Figure 1C**). Differential expression analysis using Limma package found 142 DEGs (adjusted p-value < 0.05, |log□fold change| ≥ 1.5), comprising 99 upregulated and 43 downregulated genes (**Figure 1D**). A heatmap of the top 10 upregulated (HBG2, HBD, ALAS2, PER1, SAMSN1, MALAT1, BOD1L1, CLC, LTF, and DEFA1) and top 10 downregulated (EIF5A, HINT3, LILRA5, FCAR, EIF2S3, CASP2, GM2A, ALYREF, ARF6, and SNX20) genes revealed well-differentiated expression patterns that distinguished MS patients from healthy controls with the exception of three MS samples (GSM545842, GSM545843, and GSM545845). (**Figure 1E**).

**Figure 1.**
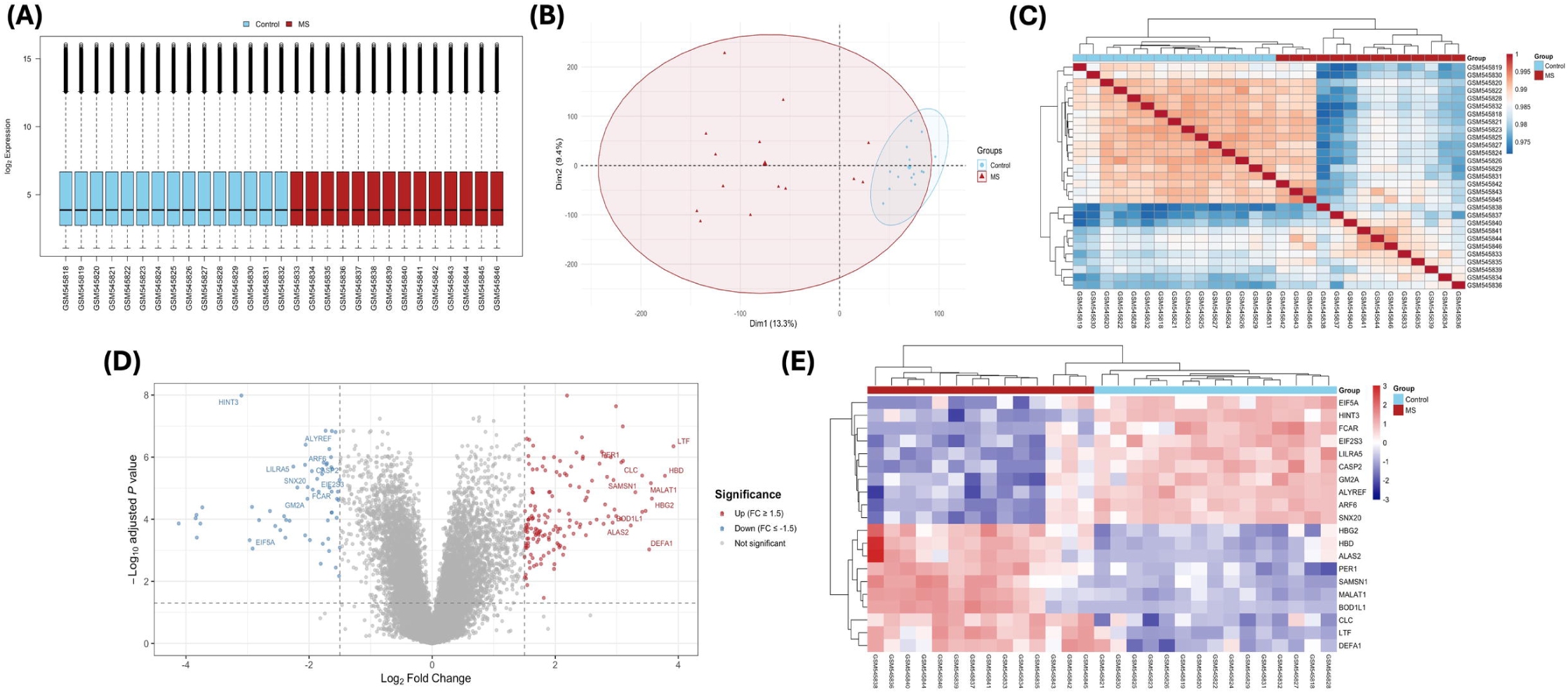
Gene expression profiling and differential expression analysis of MS vs. healthy control samples (dataset GSE21942): **(A)** Boxplots of uniform distributions of log□□transformed, quantile□normalized expression levels for 14 MS and 15 control samples. **(B)** Principal component analysis with evident discrimination between MS and control samples. **(C)** Pearson correlation heatmap with hierarchical clustering displaying evident segregation between MS and control groups. **(D)** Volcano plot of highly upregulated (red) and downregulated (blue) genes by Limma analysis (adjusted p-value < 0.05; |log□fold□change| ≥ 1.5), and the top ten in each direction. **(E)** Heatmap of different expression patterns of the top ten upregulated and top ten downregulated genes, which well distinguished MS samples from controls.

#### 3.1.2. Protein-protein Interaction Analysis, Identification of Candidate Genes and Functional Modules

STRING database was used to build the protein-protein interaction network of the 142 DEGs that were identified. The network was plotted using Cytoscape and then exported for visualization in R (**Figure 2A**). We identified the top 10 candidate genes in the network based on their Maximal Clique Centrality using the CytoHubba plugin in Cytoscape. The candidate genes were BLK, CD22, CD79A, FCRL1, FCRL5, FCRLA, MS4A1, NIBAN3, TCL1A, and VPREB3 and the network was exported to R for visualization (**Figure 2B**). Functional modules of the PPI network derived from STRING were discovered using MCODE plugin in Cytoscape and four subnetworks were obtained. Cluster 1 comprised 14 nodes with the highest MCODE score of 12.615 and was indicated as the most important functional cluster of the network (**Figure 2C**). Cluster 2 comprised 12 nodes with a score of 11.167 (**Figure 2D**). Cluster 3 contained fewer nodes with five nodes and a score of 5 (**Figure 2E**), whereas cluster 4 contained 12 nodes and the lowest score of 3.636 (**Figure 2F**). All MCODE clusters were exported from Cytoscape to R for visualization as shown. Interestingly, most genes in all four clusters were upregulated in MS patients compared to healthy controls, except for ALYREF in Cluster 3 and STAT2, CLEC7A, and ANXA1 in Cluster 4, which were downregulated.

**Figure 2.**
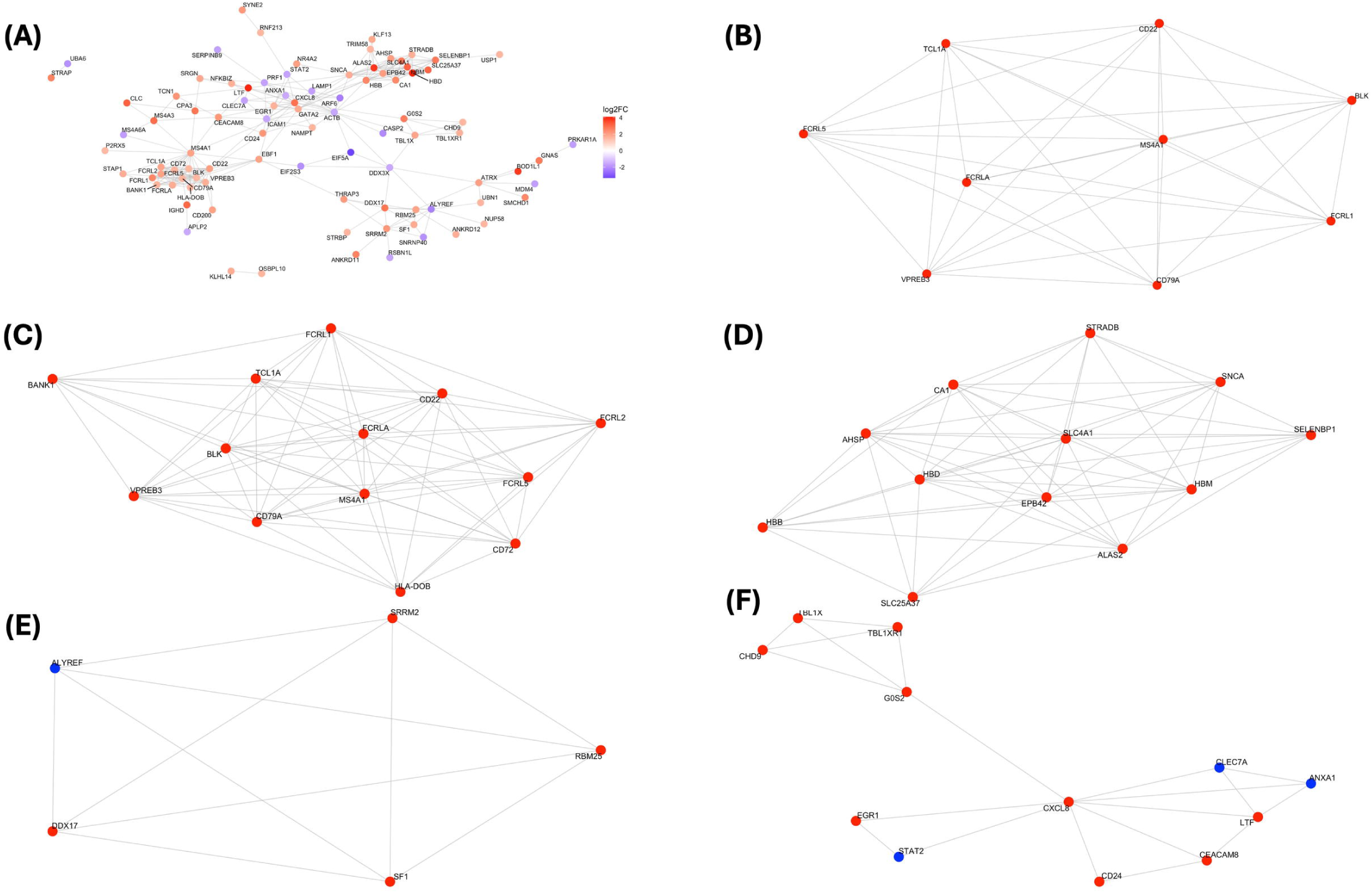
Protein-protein interaction analysis, CytoHubba candidate genes, and functional modules identification: **(A)** STRING-based protein-protein interaction network for the 142 DEGs obtained from GSE21942. **(B)** Visualization of candidate genes detected by CytoHubba according to the Maximal Clique Centrality score. **(C)** MCODE Cluster 1. **(D)** MCODE Cluster 2. **(E)** MCODE Cluster 3. **(F)** MCODE Cluster 4.

#### 3.1.3. Gene Ontology and Reactome Pathway Enrichment Analysis

To understand the biological significance of the four cluster networks obtained from the STRING-derived network we utilized the clusterProfiler and ReactomePA R packages to conduct gene ontology and reactome pathway enrichment analysis for the four clusters. Notably, Cluster 1 biological processes (BPs) were enriched for B cell activation and signal transduction pathways (**Figure 3A**). The external side of the plasma membrane and cell surface was notably overrepresented cellular compartment (CC), while signaling receptor binding and transmembrane receptor activity were notably overrepresented molecular functions (MFs). Cluster 1 Reactome pathways were antigen activation of B cell receptors and signaling by B cell receptors. Cluster 2 BPs were mainly focused on hemoglobin metabolism and oxygen and carbon dioxide transport (**Figure 3B**). The most enriched CC was hemoglobin complex, with the top MF being hemoglobin binding. The major Reactome pathway identified was erythrocyte-mediated uptake of oxygen and release of carbon dioxide. Cluster 3 reported highly significant enrichment for RNA splicing and mRNA processing among BPs, of which the most enriched CC was spliceosome and the most enriched MF was mRNA binding (**Figure 3C**). Expectedly, the most enriched Reactome pathway was mRNA splicing. Cluster 4 was primarily associated with immune system activation (**Figure 3D**). DNA binding was emphasized as the primary MF, and the RAR-related orphan receptor alpha activation pathway was the most strongly enriched Reactome pathway.

**Figure 3.**
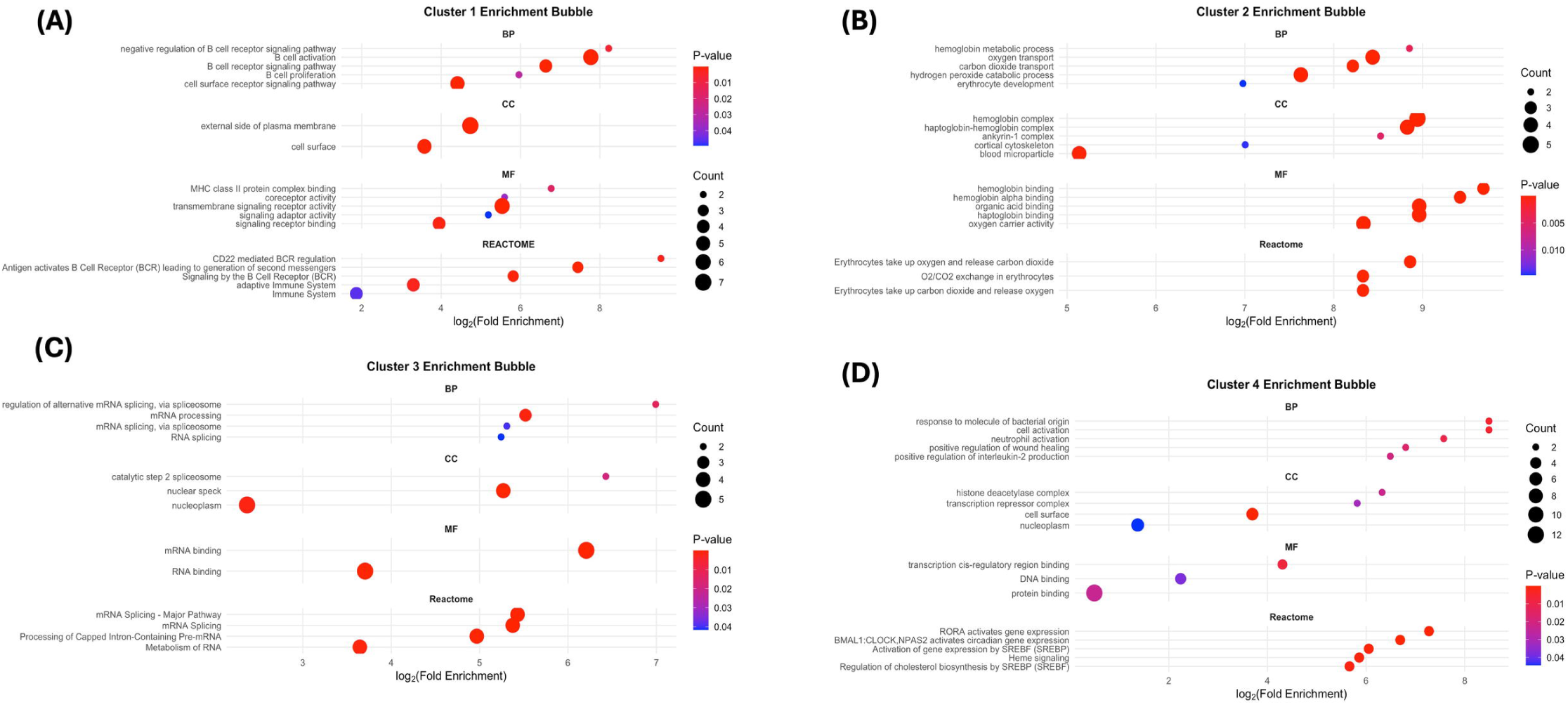
Gene ontology and reactome pathway analysis: **(A)** Cluster 1 enrichment bubble. **(B)** Cluster 2 enrichment bubble. **(C)** Cluster 3 enrichment bubble. **(D)** Cluster 4 enrichment bubble.

#### 3.1.4. Weighted Gene Co-expression Network Analysis

WGCNA of the PBMCs transcriptome segregated expressed genes into more than 30 discrete clusters as shown in the dendrogram in (**Figure 4A**). Network topology analysis of the scale-free topology (left sub-panel) indicates that the fit index (signed R²) increases hugely with increasing soft-thresholding parameter β, first breaking the conventional 0.80 threshold at β = 4. Meanwhile, mean connectivity (right sub-panel) decreases dramatically from approximately 3500 at β = 1 to nearly zero by β = 6 (**Figure 4B**). Therefore, we chose a soft-thresholding power 4 as it creates a network that is both scale-free and connected sufficiently for downstream analysis. The eigengene correlation heat-map with disease status identified seven modules that were significantly correlated with MS disease status. The most significant clusters identified were two upregulated modules (blue and purple) and two downregulated modules (turquoise and brown) (**Figure 4C**). In particular, the blue and purple eigengenes are significantly similarly correlated with MS samples (upregulated in MS), and the turquoise and brown eigengenes are significantly oppositely correlated with the disease (downregulated in MS). For each of the four disease-associated modules, the ten most highly connected intramodular hub genes are displayed (**Figure 4D**). In total, the profile identifies two induced transcriptional programs in MS PBMCs (blue and purple) and two repressed programs (turquoise and brown).

**Figure 4.**
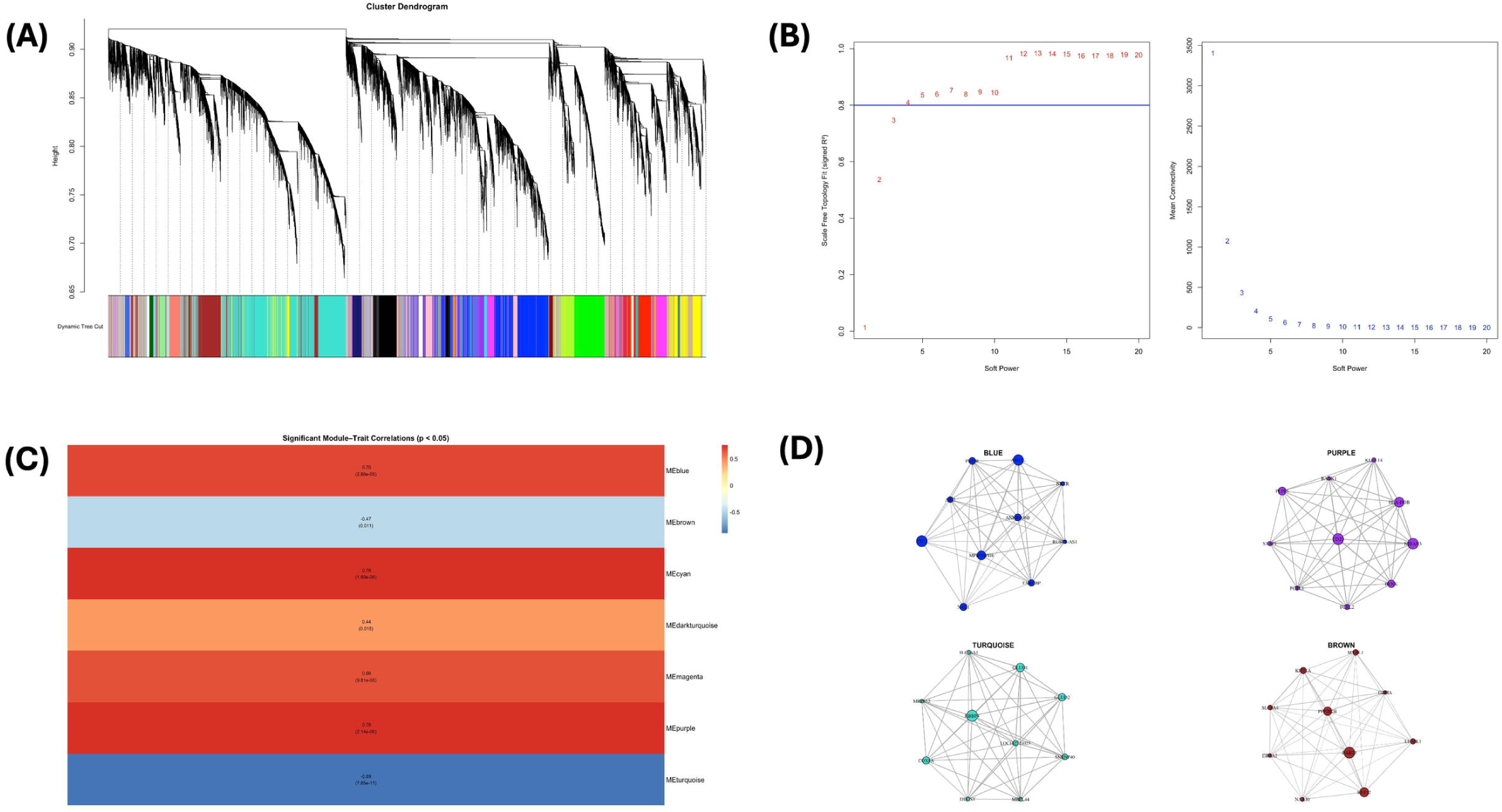
WGCNA: **(A)** WGCNA gene cluster dendrogram showing module assignment. **(B)** Scale independence and mean connectivity across soft-thresholding powers. **(C)** Module-trait relationships heat map with warm colors denoting positive correlation and cool colors negative correlation with disease. **(D)** Top 10 gene networks of significant modules that correlate with disease status.

#### 3.1.5. WGCNA Cluster Modules Functional Enrichment

To probe their biological roles, we subjected the four disease-associated modules identified by WGCNA to Gene Ontology-Biological Process enrichment analysis. The upregulated blue module is enriched for RNA splicing and ribosome biogenesis functions (**Figure 5A**), whereas the upregulated purple module is enriched for B-cell activation and antigen-receptor signaling pathways (**Figure 5B**). In contrast, the downregulated turquoise module is enriched for mitochondrial respiration and oxidative phosphorylation processes (**Figure 5C**), and the downregulated brown module is enriched for natural-killer/T-cell–mediated cytotoxicity and lymphocyte-differentiation pathways (**Figure 5D**).

**Figure 5.**
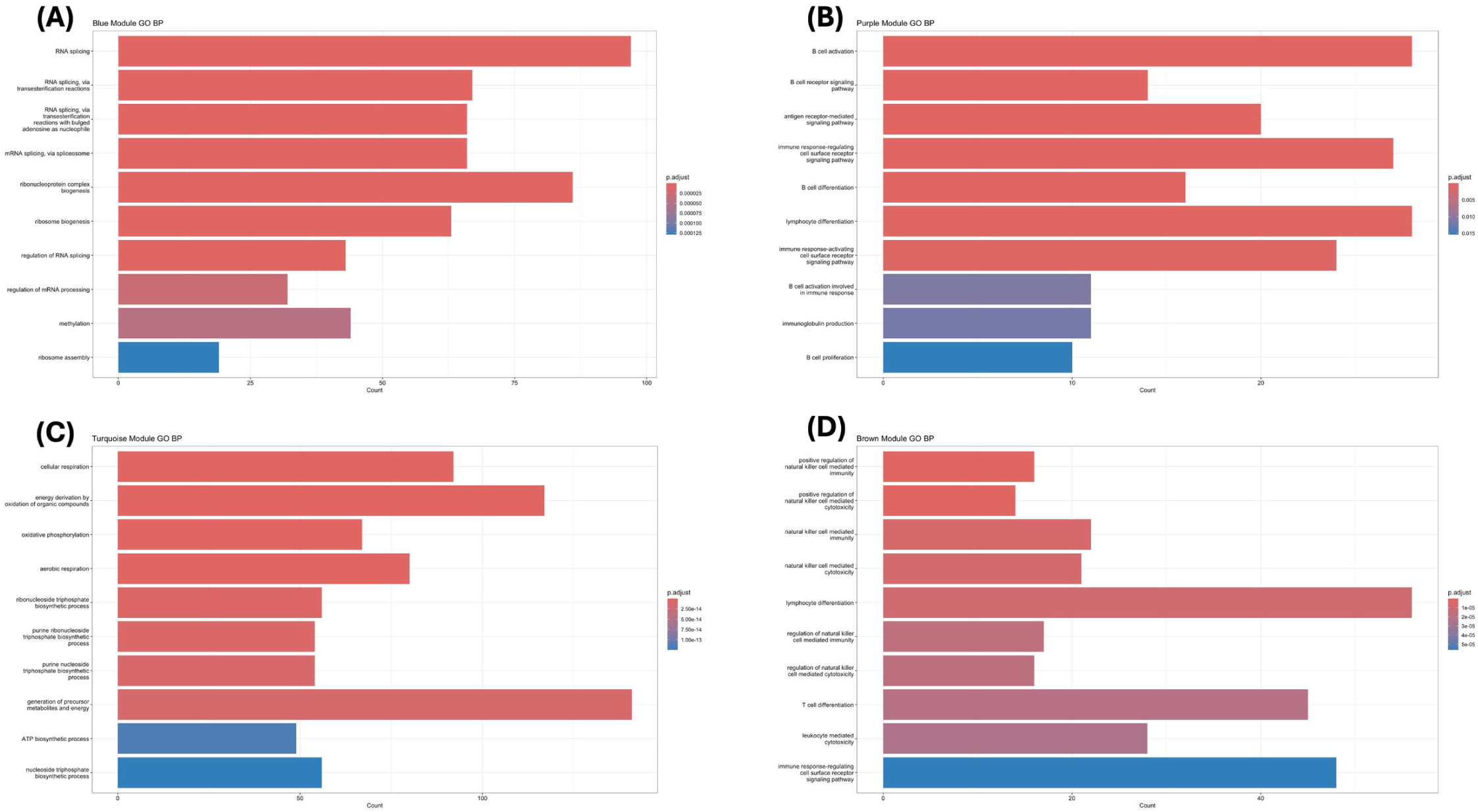
Gene Ontology–Biological Process enrichment of the four disease-associated WGCNA modules. **(A)** Blue module. **(B)** Purple module. **(C)** Turquoise module. **(D)** Brown module.

#### 3.1.6. Hub Genes Identification

To identify the hub genes from candidate genes we used LASSO machine learning algorithm with stratified five-fold cross validation. The cross-validation tuning curve (**Figure 6A**) achieved its highest mean AUC at λ_min_ , with the “1 SE” rule selecting a slightly larger λ that yielded a three-gene model (FCRL1, CD22, and CD79A). As λ decreases, gene coefficients remain at zero under strong regularization, then begin to diverge from zero as the penalty weakens. By the time we reach λ□□□, only FCRL1, CD22, and CD79A have non-zero coefficients (**Figure 6B**). The final three□gene model achieved an AUC of 0.957 and 0.85 accuracy with a 0.88 sensitivity and 0.83 specificity, indicating excellent discrimination (**Figure 6C**). Nested cross-validation confirmed the findings with an AUC of 0.978, demonstrating robust discriminatory power of FCRL1, CD22, and CD79A.

**Figure 6.**
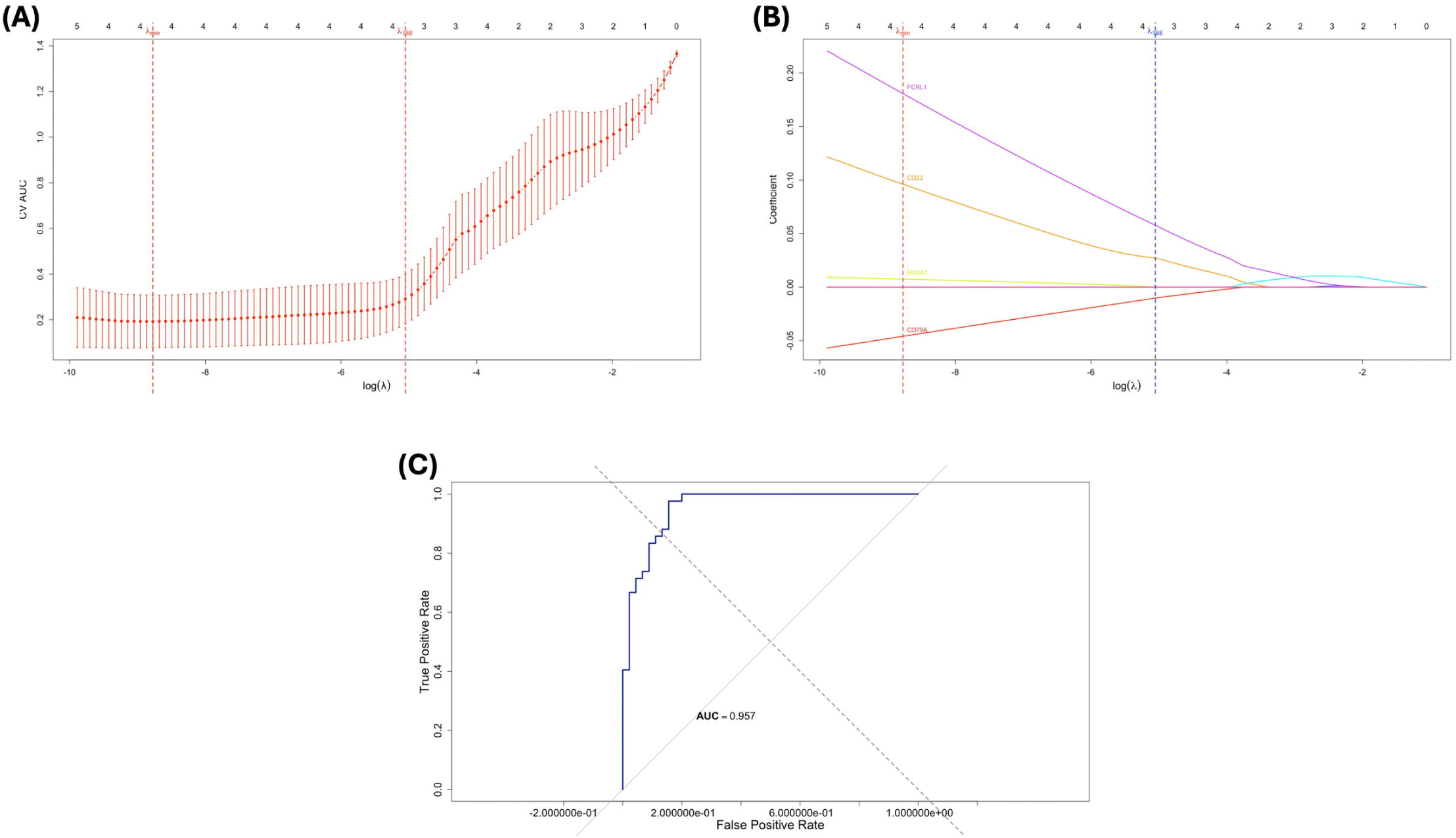
PBMCs LASSO regression analysis: **(A)** Mean cross-validated AUC plotted against log λ during stratified five-fold LASSO tuning. **(B)** Genes LASSO coefficient profiles across lambda values. **(C)** ROC curve for the final three gene LASSO model combing predictions from all five cross validation folds.

### 3.2. Brain

#### 3.2.1. Identification of Brain DEGs

Gene expression analysis on the GSE38010 dataset was performed to obtain DEGs between MS lesions and normal control brain tissue. After log□ transformation and quantile normalization, the expression profiles exhibited uniformly consistent distributions in all samples with no statistically significant outliers, confirming preprocessing integrity (**Figure 7A**). Principal component analysis revealed a partial segregation of controls and MS samples, with controls more clumped and MS samples having intra-group heterogeneity that could be due to heterogeneity of lesions (active plaque, chronic activate plaque, and chronic plaque) (**Figure 7B**). This was supported by the correlation heatmap, where Pearson correlation coefficients were modestly reduced compared to GSE21942 with a minimum correlation value approaching 0.7, but still remained able to display grouping by disease status (**Figure 7C**). Differential expression analysis picked 1,493 DEGs between MS and control groups with Limma (adjusted p-value < 0.05 and |log□ fold-change| ≥ 1.5), with nearly an equal number of upregulated and downregulated DEGs (**Figure 7D**). Top 10 upregulated genes (UNC13C, ITPKA, GPR22, NWD2, AKAP5, RXFP1, CELF4, PRR36, C3orf80, and TRHDE) and top 10 downregulated genes (LIPF, SCD, FGFR2, RHBDL2, DLK1, S1PR5, HSD3B2, MBP, CYP17A1, and TTR) heatmap exhibited moderate but clear transcriptional heterogeneity among MS lesion subtypes and control tissues (**Figure 6E**), irrespective of expression level variation among individual lesions.

**Figure 7.**
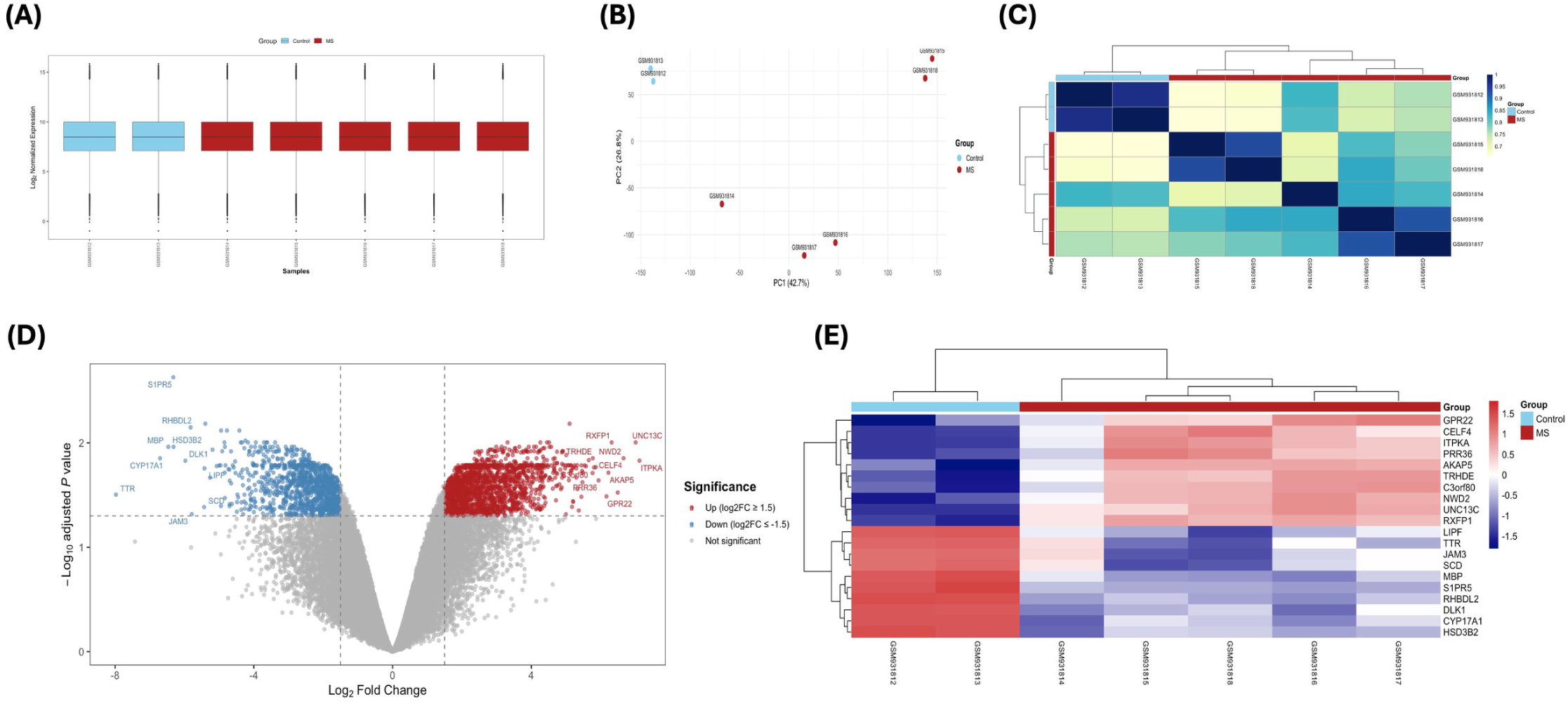
Gene expression profiling and differential expression analysis of MS vs. control brain samples (dataset GSE38010): **(A)** Boxplots of uniform distributions of log□□transformed, quantile□normalized expression levels for 5 MS and 2 control samples. **(B)** Principal component analysis with evident discrimination between MS and control samples. **(C)** Pearson correlation heatmap with hierarchical clustering displaying evident segregation between MS and control groups. **(D)** Volcano plot of highly upregulated (red) and downregulated (blue) genes by Limma analysis (adjusted p-value < 0.05; |log□fold□change| ≥ 1.5), and the top ten in each direction. **(E)** Heatmap of different expression patterns of the top ten upregulated and top ten downregulated genes, which well distinguished MS samples from controls.

#### 3.2.2. Protein-Protein Interaction Analysis, Identification of Candidate Genes and Functional Modules

STRING database was used to build the protein-protein interaction network of the 1,493 DEGs that were identified. The network was plotted using Cytoscape and then the top 500 DEGs in terms of log FC was exported for visualization in R and the top 10 upregulated and top 10 downregulated genes were named in the network (**Figure 8A**). We identified the top 10 candidate genes in the network based on their Maximal Clique Centrality using the CytoHubba plugin in Cytoscape. The candidate genes were DLGAP1, DLGAP2, GRIA1, GRIA2, GRIN1, GRIN2A, HOMER1, NLGN1, NRXN1, and SHANK2 and the network was exported to R for visualization (**Figure 8B**). Functional modules of the PPI network derived from STRING were discovered using MCODE plugin in Cytoscape and four subnetworks were obtained. Cluster 1 comprised 50 nodes with the highest MCODE score of 14.122 and was indicated as the most important functional cluster of the network (**Figure 8C**). Cluster 2 comprised 56 nodes with a score of 6.182 (**Figure 8D**). Cluster 3 contained fewer nodes with five nodes and a score of 4.5 (**Figure 8E**), whereas cluster 4 contained 70 nodes and the lowest score of 4.377 (**Figure 8F**). All MCODE clusters were exported from Cytoscape to R for visualization as shown.

**Figure 8.**
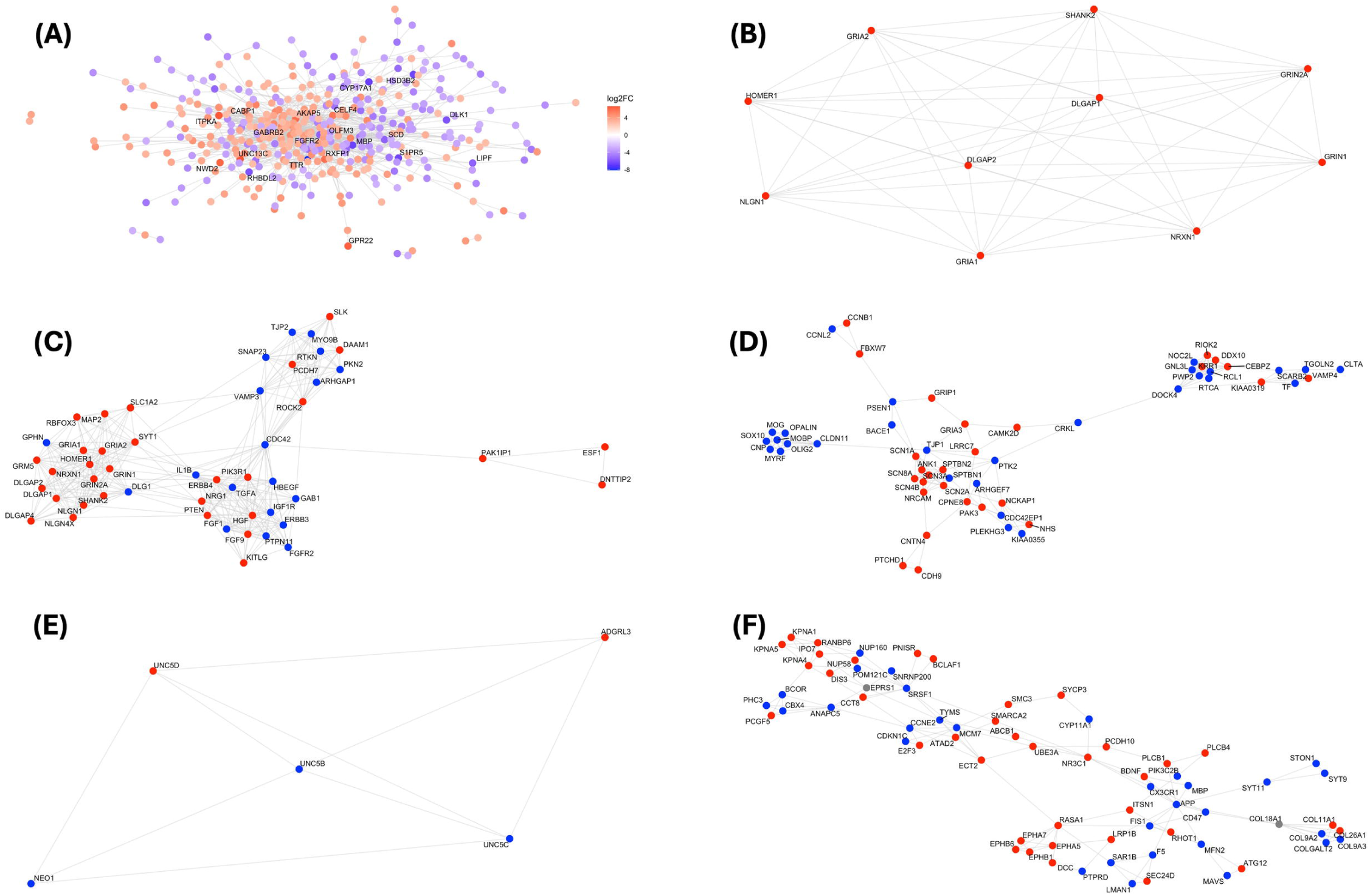
Protein-protein interaction analysis, CytoHubba candidate genes, and functional modules identification: **(A)** STRING-based protein-protein interaction network for the 1,493 DEGs obtained from GSE38010. **(B)** Visualization of candidate genes detected by CytoHubba according to the Maximal Clique Centrality score. **(C)** MCODE Cluster 1. **(D)** MCODE Cluster 2. **(E)** MCODE Cluster 3. **(F)** MCODE Cluster 4.

#### 3.2.3. Gene Ontology

To understand the biological significance of the four cluster networks obtained from the STRING-derived network we utilized the clusterProfiler R package to conduct gene ontology for the four clusters. Cluster 1 was enriched with chemical synaptic transmission and positive regulation of excitatory postsynaptic potentials, with glutamatergic synapse being one of the CCs (**Figure 9A**). Cluster 2 was enriched in axon and neuronal ensheathment processes and the voltage-gated sodium channel complex was the top CC and voltage-gated sodium channel activity was the top MF (**Figure 9B**). For Cluster 3, netrin-activated signal pathways were the top BP and netrin receptor activity was the top highly enriched MF (**Figure 9C**). The most enriched BP in Cluster 4 was protein import into the nucleus, and the most enriched CC and MF were nuclear pore and nuclear localization sequence binding, respectively (**Figure 9D**).

**Figure 9.**
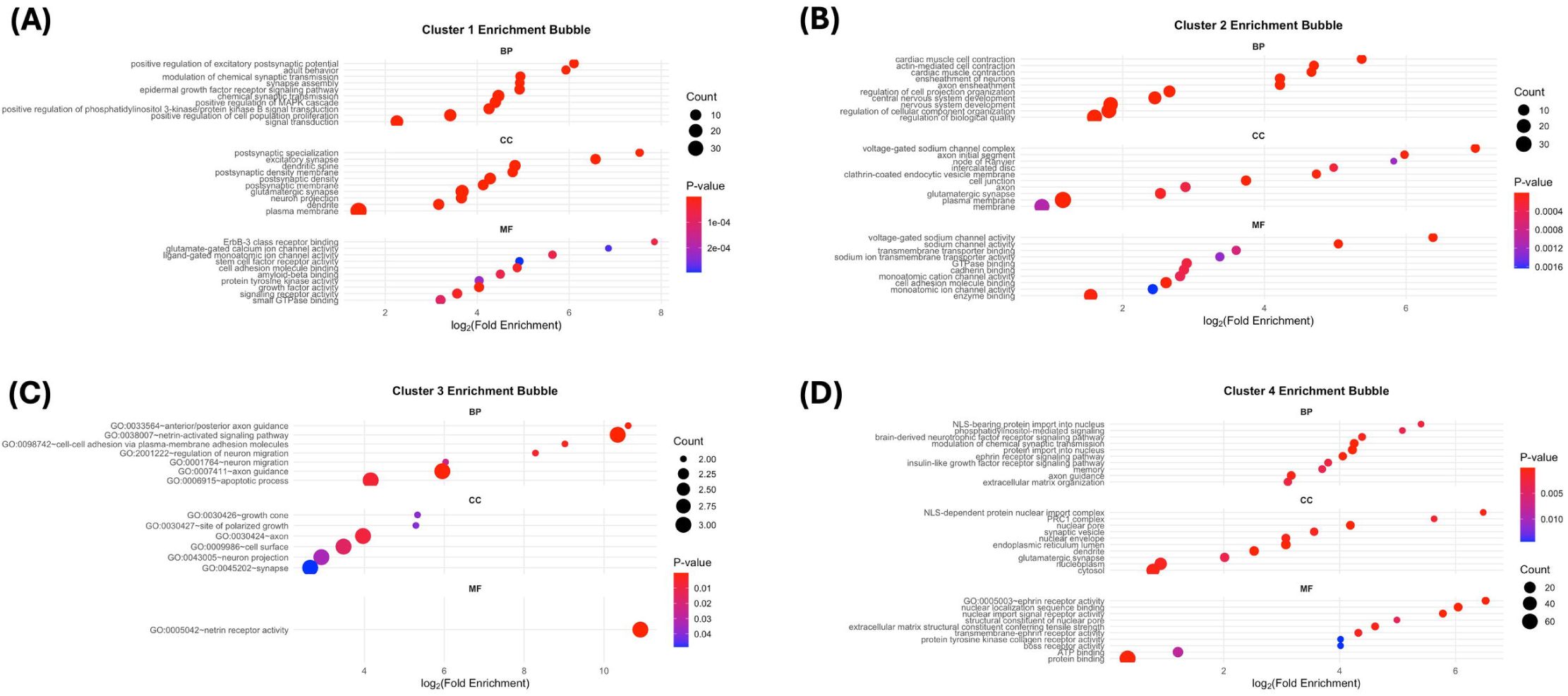
Gene ontology: **(A)** Cluster 1 enrichment bubble. **(B)** Cluster 2 enrichment bubble. **(C)** Cluster 3 enrichment bubble. **(D)** Cluster 4 enrichment bubble.

#### 3.2.4. Gene Set Enrichment Analysis

GSEA was used to delineate the upregulated and downregulated biological processes, reactome pathways, and WikiPathways within the identified DEGs. Synaptic signaling, regulation of postsynaptic membrane potential, and modulation of chemical synaptic transmission were upregulated biological processes identified by GSEA (**Figure 10A**). Ensheathment of axons and neurons, oligodendrocyte differentiation, and steroid metabolic processes were downregulated biological processes (**Figure 10B**). The upregulated reactome pathways were neurotransmitter receptors and postsynaptic signal transmission and transmission across chemical synapse (**Figure 10C**). Downregulated reactome pathways were involved mainly in steroid and lipid metabolism (**Figure 10D**). Neuroinflammation and glutamatergic signaling was an upregulated WikiPathway (**Figure 10E**) and the downregulated WikiPathways are shown in (**Figure 10F**).

**Figure 10.**
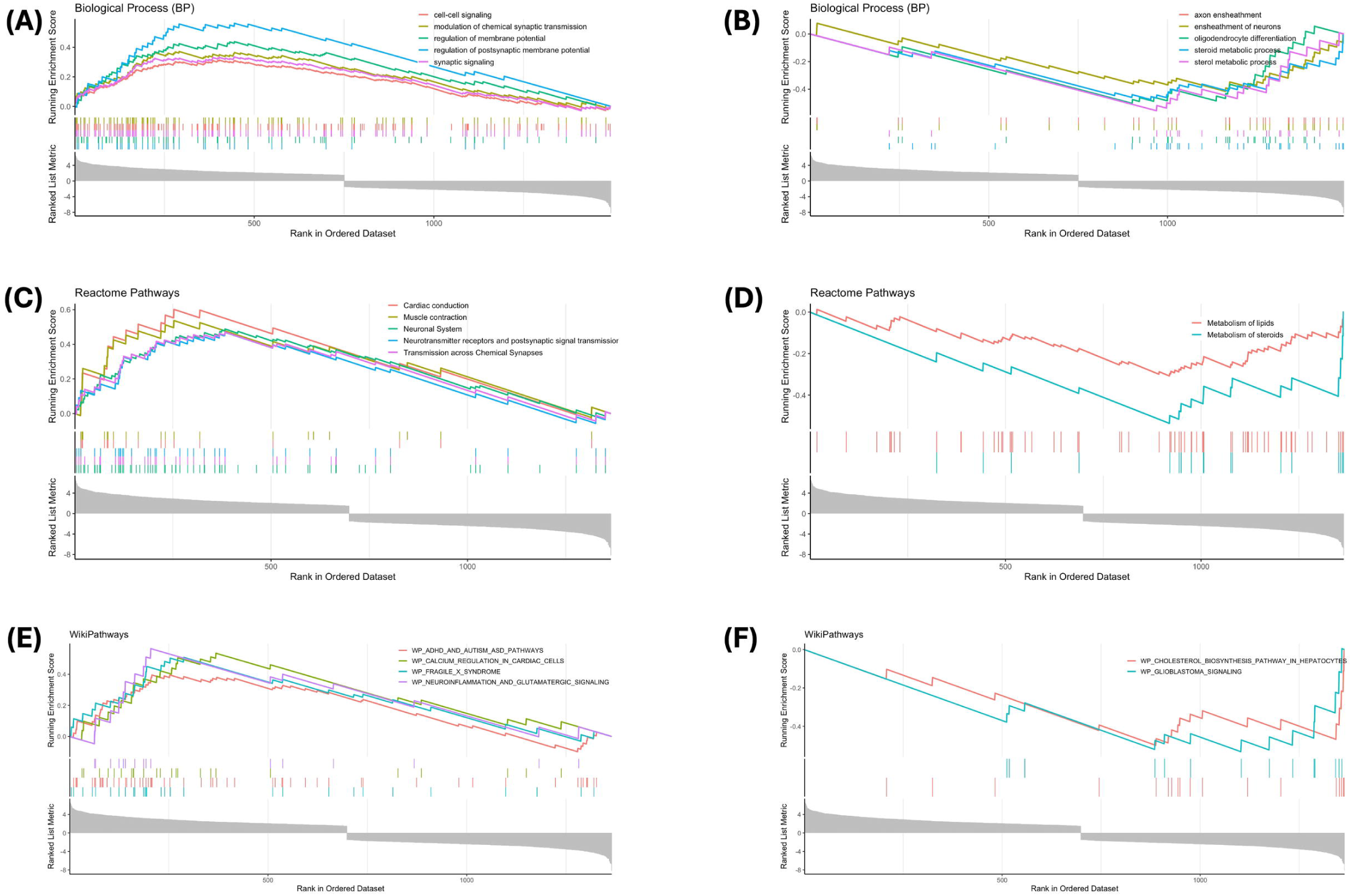
GSEA: **(A)** Upregulated BPs. **(B)** Downregulated BPs. **(C)** Upregulated Reactome pathways. **(D)** Downregulated Reactome pathways. **(E)** Upregulated WikiPathways. **(F)** Downregulated WikiPathways.

### 3.3. Molecular Docking Study

#### 3.3.1. Virtual Screening Analysis

In order to find out the therapeutic leads for our selected hub genes, we tested a library of Selleckchem ligands in a molecular docking study against four hub genes CD79A, GRIN2A, NRXN1, and GRIA2. We chose only those compounds with high BBB permeability, good bioavailability scores, and not violating Lipinski’s, Ghose’s, Veber’s, Egan’s, and Muegge’s rules as shown in the supplementary data section. The 20 best compounds that interact with both targets, sorted based on binding energy, are listed in supplementary data. Icotinib and Niraparib were two of the highest affinity compounds to four targets. Icotinib showed -6.3 Kcal/mol binding energy against CD79A, -9.8 Kcal/mol binding energy against GRIN2A, -7.1 Kcal/mol binding energy against NRXN1, and -8.2 Kcal/mol binding energy against GRIA2, and Niraparib showed -6.6 Kcal/mol binding energy against CD79A, -9.4 Kcal/mol binding energy against GRIN2A, -7 Kcal/mol binding energy against NRXN1, and -7.9 Kcal/mol binding energy against GRIA2, revealing these drugs as promising candidates for further investigation. However, based on our MD simulations only CD79A and GRIA2 produced satisfactory results with both icotinib and niraparib and their molecular docking, dynamics, and free energy analysis are highlighted below. Molecular docking and dynamics results for GRIN2A and NRXN1 are in the supplementary materials ( Figures S1, S2, S3, and S4).

#### 3.3.2. Molecular docking of icotinib and niraparib with CD79A

Molecular docking simulation unveiled the binding interaction between icotinib and niraparib to CD79A. Both ligands occupied a common binding site on the CD79A protein where icotinib (green) and Niraparib (red) had energetically favorable conformations inside the site (**Figure 11A**). High-resolution 2D interaction maps revealed that icotinib established varied van der Waals interactions, normal and carbon hydrogen bonds, π-π stacked, and π-σ interaction with important residues like ARG68, GLU79, LEU70, GLY102, and TRP76 (**Figure 11B**). Niraparib also established interactions with a number of important residues like ARG68, VAL69, LEU70, and PRO77 with van der Waals forces, π-cation and π-σ contacts, and conventional hydrogen bonds (**Figure 11C**). Together, these interactions highlight key residues mediating ligand recognition and support the binding potential of both compounds toward CD79A.

**Figure 11.**
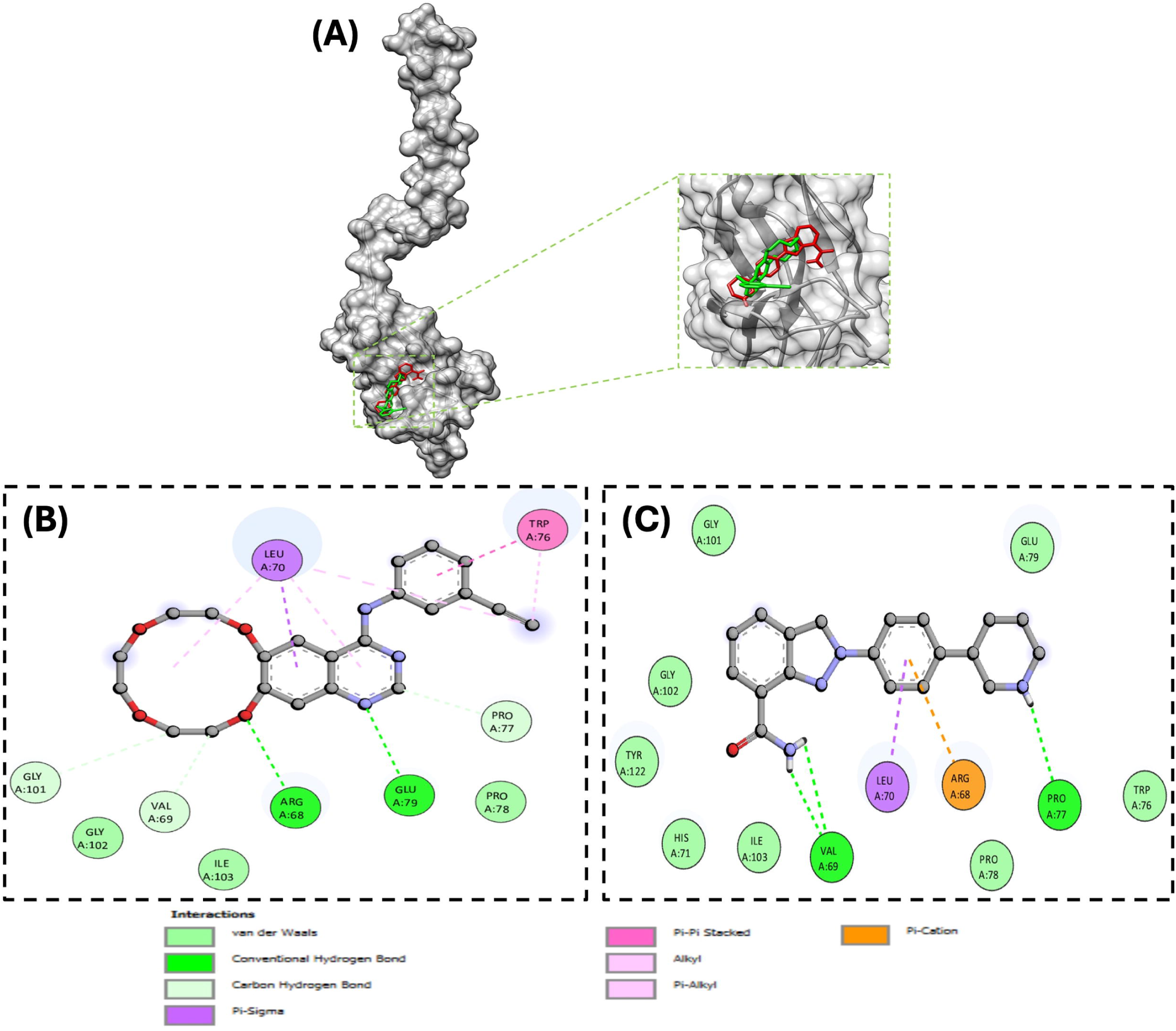
Molecular docking of icotinib and niraparib with CD79A: **(A)** The 3D binding configuration of icotinib (Green) and niraparib (Red). The molecular interaction fingerprints in 2D showed that atoms of **(B)** icotinib and **(C)** niraparib interacted with amino acid residues important for binding with CD79A.

#### 3.3.3. Molecular docking of icotinib and niraparib with GRIA2

Molecular docking study demonstrated the binding interaction of icotinib and niraparib with GRIA2. Both ligands were docked inside the same active site of GRIA2 where Icotinib (green) and Niraparib (red) filled overlapping areas of the binding pocket (**Figure 12A**). The 2D interaction plots also gave a clear impression of the character of ligand–protein interactions. Icotinib made diverse stabilizing interactions, including traditional hydrogen bonds with ARG96 and THR91, and a π-anion interaction with GLU193. There were other van der Waals and π–π stacked interactions with the likes of TYR220 and TYR61 (**Figure 12B**). Niraparib engages the same pocket, forming traditional hydrogen bonding with ARG96 and GLY62 and van der Waals interactions with residues such as SER142, GLU193, and MET196, and additional hydrophobic/ π-type interactions with TYR61, THR93, PRO89, and TYR220 (**Figure 12C**). Together, these interactions highlight key residues mediating ligand recognition and support the binding potential of both compounds toward GRIA2.

**Figure 12.**
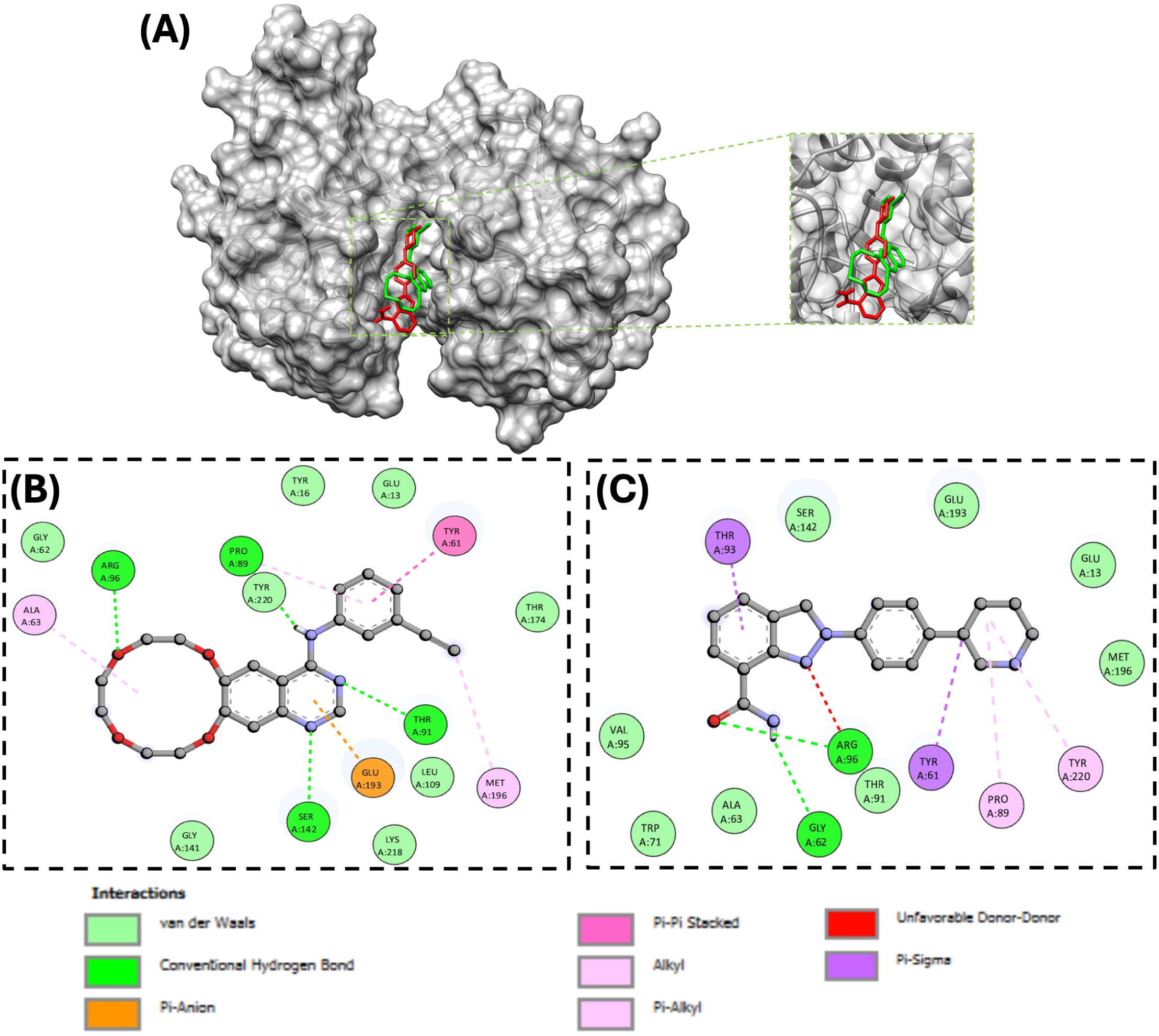
Molecular docking of icotinib and niraparib with GRIA2: **(A)** The 3D binding configuration of icotinib (Green) and niraparib (Red). The molecular interaction fingerprints in 2D showed that atoms of **(B)** icotinib and **(C)** niraparib interacted with amino acid residues important for binding with GRIA2.

#### 3.3.4. Molecular dynamics simulation of icotinib and niraparib with CD79A

Molecular-dynamics simulation analysis of icotinib (blue) and niraparib (red) bound to CD79A on a 200 ns trajectory is presented. RMSD trace indicates that both complexes equilibrate within 20 ns; the niraparib–CD79A complex (red) then stabilizes to a lower plateau i.e., greater backbone stability, while the Icotinib complex (blue) oscillates in a rather narrower range around a higher baseline (**Figure 13A**). A comparable pattern is observed in the RMSF representation of icotinib with an overall lower residue-level motions compared to niraparib, and the niraparib complex is slightly more dynamic (**Figure 13B**). The radius of gyration is little reduced for niraparib, implying a slightly more compact overall structure (**Figure 13C**). This tightness is also evident in the solvent-accessible surface area, where niraparib has lower SASA compared to icotinib, reflecting better-buried protein surface (**Figure 13D**). Finally, hydrogen-bond analysis shows that icotinib has the highest average number of bonds especially in the second half of the simulation reflecting higher and more stable network of contacts with CD79A (**Figure 13E**). Overall, niraparib gives a lower backbone RMSD, while icotinib compensates with a stronger hydrogen-bond network in the late stage and less fluctuations, making the two ligands both good but highlighting different features of stability.

**Figure 13.**
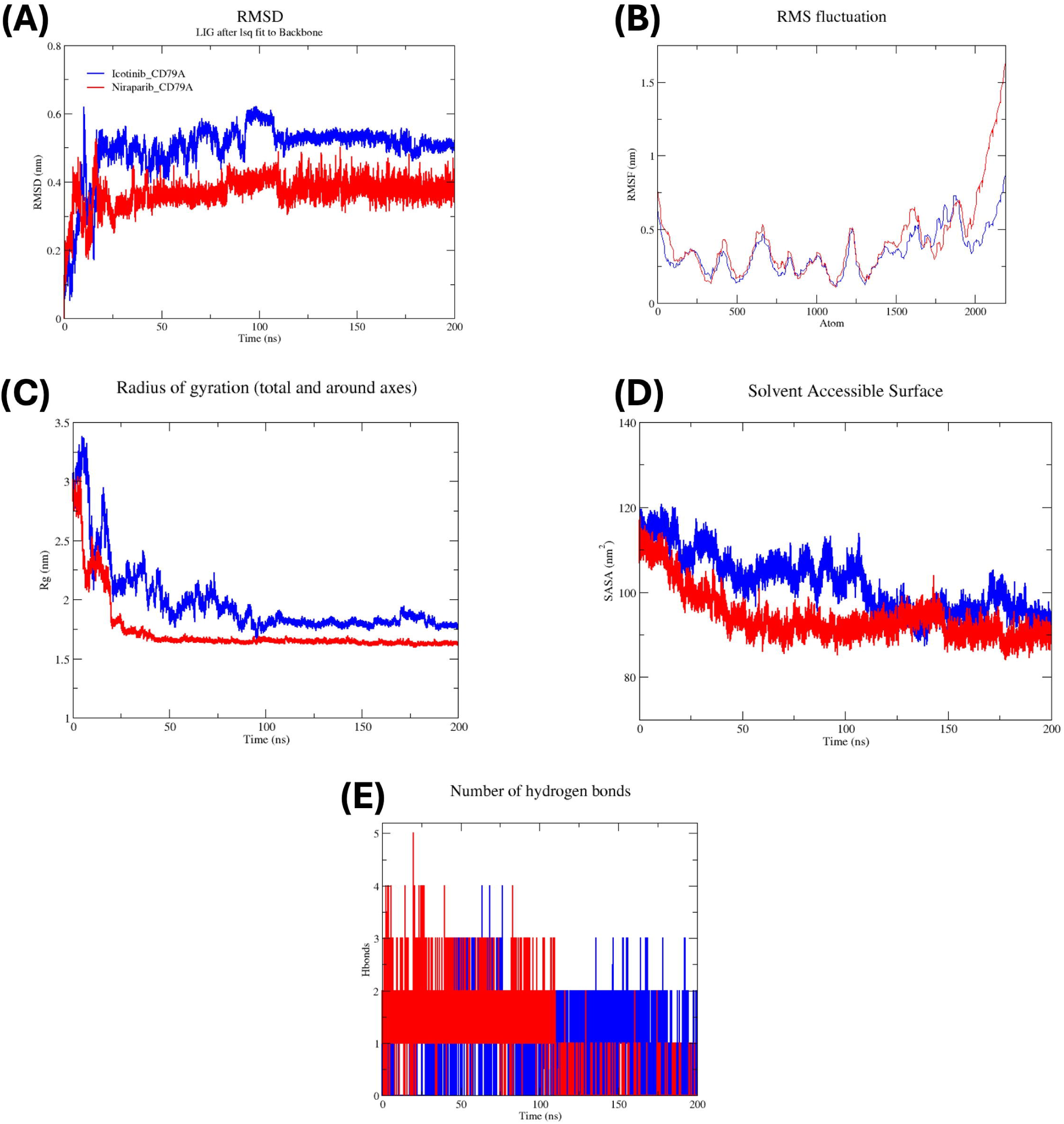
Molecular dynamics simulation analysis of icotinib and niraparib complexes with CD79A: **(A)** RMSD, **(B)** RMSF, **(C)** Radius of Gyration, **(D)** Solvent Accessible Surface Area, and **(E)** Hydrogen Bond Profiles Over 200 ns Simulations.

#### 3.3.5. Molecular dynamics simulation of icotinib and niraparib with GRIA2

The molecular-dynamics simulation trajectory of the GRIA2 complex with icotinib (cyan) and niraparib (magenta) for 200 ns is presented. The RMSD traces indicate that both complexes become stabilized within the initial ∼20 ns; however, the niraparib–GRIA2 complex (magenta) has reduced deviations and plateaus at a lower RMSD thereafter, reflecting higher global stability (**Figure 14A**). This tendency is replicated within the RMSF profile, where niraparib creates smaller residue-level fluctuations, whereas icotinib shows increased mobility (**Figure 14B**). In the radius-of-gyration plot icotinib starts out with a more open arrangement but increasingly becomes more closed as the run goes on, ending up more closed than niraparib’s complex (**Figure 14C**). The solvent-accessible surface area is similarly larger to start with for icotinib, then gradually decreasing down until it falls below the niraparib profile showing progressive burial of the protein surface (**Figure 14D**). Hydrogen-bond analysis reveals that niraparib maintains a higher and longer-term hydrogen-bond numbers along most of the path, which suggests a tighter interaction network (**Figure 14E**). In sum, niraparib delivers the steadier, better-engaged complex, while icotinib ultimately leaves GRIA2 in a more compact and solvent-shielded state.

**Figure 14.**
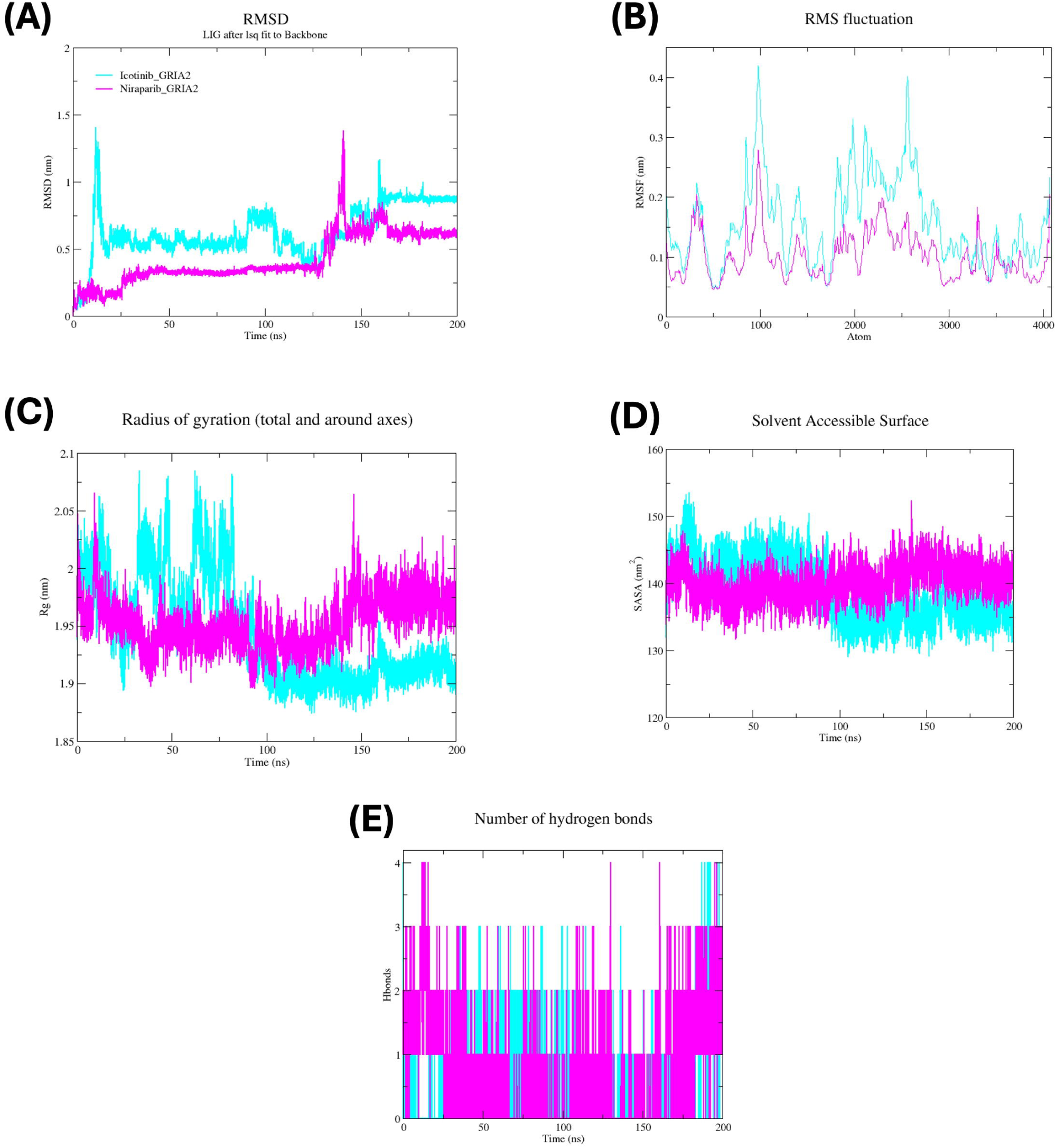
Molecular dynamics simulation analysis of icotinib and niraparib complexes with GRIA2: **(A)** RMSD, **(B)** RMSF, **(C)** Radius of Gyration, **(D)** Solvent Accessible Surface Area, and **(E)** Hydrogen Bond Profiles Over 200 ns Simulations.

#### 3.3.6. Binding free energy calculation of icotinib and niraparib with CD79A

Figure 15 illustrates the MM/PBSA binding free-energy decomposition for icotinib (A) and niraparib (B) in complex with CD79A. In the inset panels, icotinib binding is marked by a favorable van der Waals interaction with smaller but still favorable electrostatic and non-polar solvation terms; these are offset to some extent by a disfavorable polar solvation penalty, resulting in a net negative (favorable) ΔG (**Figure 15A, left**). Niraparib possesses an extremely large favorable electrostatic contribution and a favorable van der Waals term but are largely counterbalanced by a large polar solvation penalty, and the net ΔG is still favorable but less negatively sloped than icotinib (**Figure 15C, left**). In the per-residue decomposition, the ligand term (B:LIG:172) provides the dominant stabilizing contribution in each complex. Among the protein residues: LEU70, TRP76, and PRO78 provide stabilizing energy towards icotinib (**Figure 15A, right**), while LEU70, TRP76, GLU79, VAL69, and GLY101 facilitate niraparib binding (**Figure 15C, right**). In addition, heat maps of the per-residue decomposition of icotinib (**Figure 15B**) and niraparib (**Figure 15D**) are shown. Collectively, the energy components indicate dispersion-dominated stabilization for Icotinib and electrostatics-enhanced (but desolvation-penalized) stabilization for niraparib.

**Figure 15.**
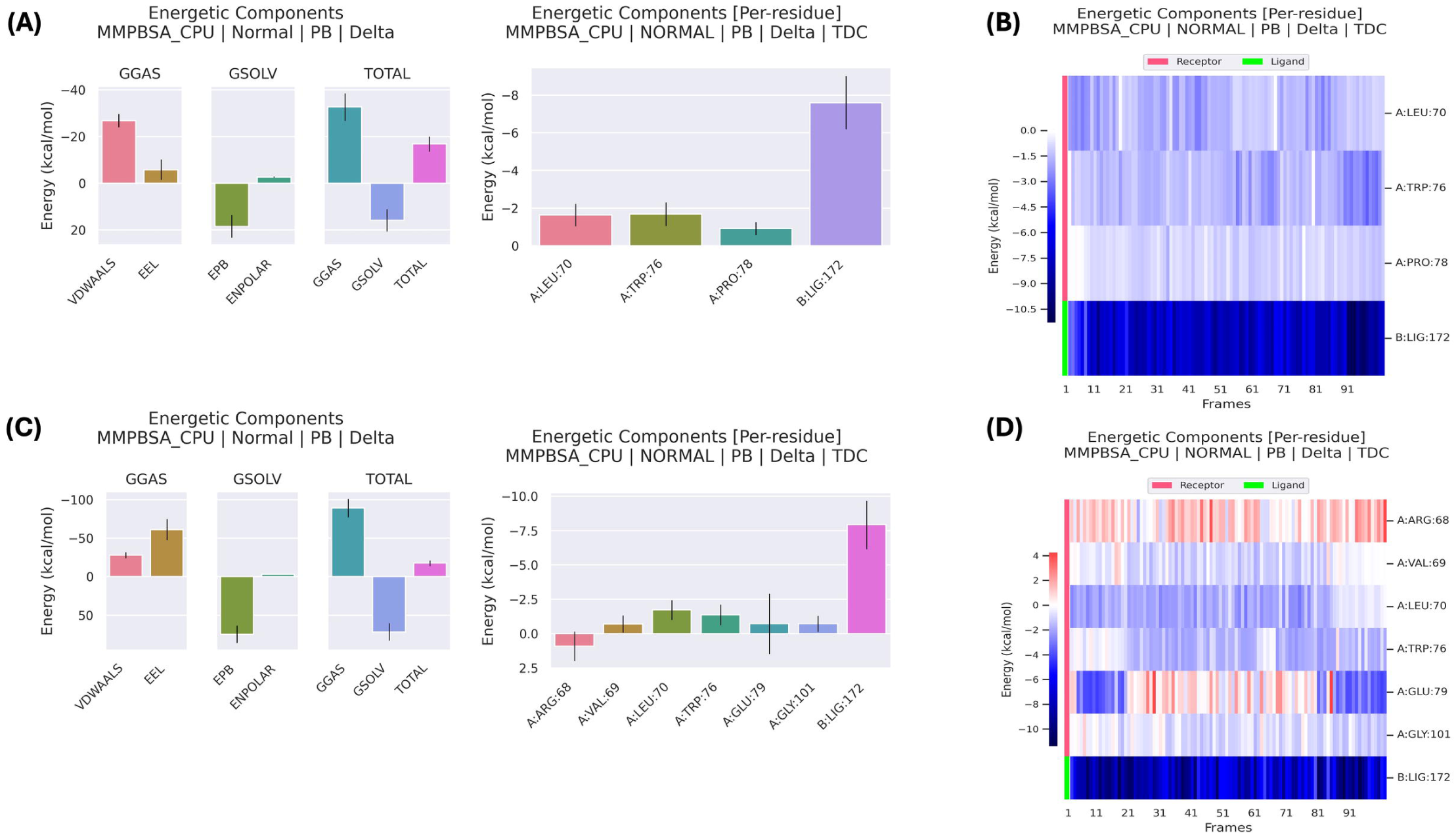
Binding free energy for icotinib and niraparib with CD79A: **(left)** binding free energy contribution by various interactions, **(right)** binding free energy contribution by active residues and ligand. **(A)** icotinib, and **(C)** niraparib. **(B)** Heat map of the per-residue decomposition of icotinib with CD79A. **(D)** Heat map of per-residue decomposition of niraparib with CD79A.

#### 3.3.7. Binding free energy calculation of icotinib and niraparib with GRIA2

Figure 16 illustrates the MM/PBSA binding free-energy decomposition of icotinib (A) and niraparib (B) bound to GRIA2. Icotinib binding is characterized in inset panels by dominant positive van der Waals and minor electrostatic and non-polar solvation terms combined to cover a broad polar solvation penalty (**Figure 16A, left**). Niraparib is stabilized by van der Waals interactions as well, whereas the electrostatic contribution is close to zero and polar desolvation penalty is less (**Figure 16C, left**); total ΔG is favorable for both the ligands with niraparib slightly higher. Decomposition per-residue indicates that the majority of stabilizing energy comes from the ligand itself (B:LIG:262) in both systems. In the protein residues, THR91 and MET196 contribute favorably to icotinib, while GLU193 is the primary favorable contributor to niraparib (**Figure 16A,C, right**). In addition, heat maps of the per-residue decomposition of icotinib (**Figure 16B**) and niraparib (**Figure 16D**) Thus, GRIA2 binding is strongly ligand-directed with residue-specific stabilization being less prominent.

**Figure 16.**
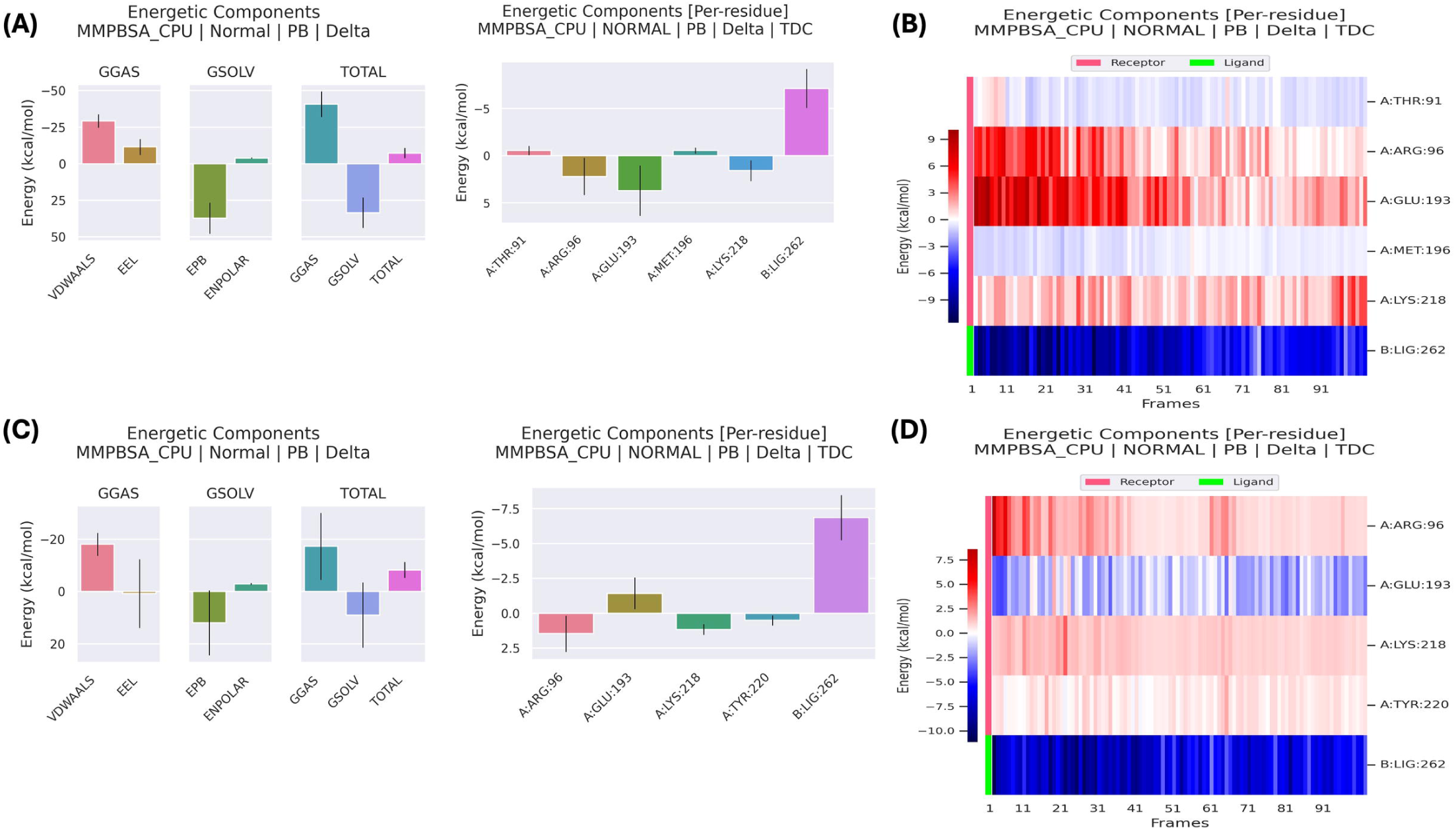
Binding free energy for icotinib and niraparib with GRIA2: (**left**) binding free energy contribution by various interactions, (**right**) binding free energy contribution by active residues and ligand. **(A)** icotinib, and **(C)** niraparib. **(B)** Heat map of the per-residue decomposition of icotinib with GRIA2. **(D)** Heat map of per-residue decomposition of niraparib with GRIA2.

## 4. Discussion

MS originally was thought to be primarily a T-cell–driven autoimmune encephalomyelitis wherein peripherally originating lymphocytes penetrate the BBB and induce focal demyelination [41]; yet two decades of therapeutic experience demonstrate that even highly selective lymphocyte-depleting regimens retards but does not completely arrest MS progression. Coles et al., reported that nearly a quarter of patients treated by the lymphocyte depleting agent alemtuzumab accumulated six month confirmed disability worsening, despite sustained suppression of relapses and absence of new magnetic resonance imaging activity [42]. Maggi et al., demonstrated that anti-CD20 depletion of B-cells suppresses new inflammatory activity yet leaves chronic active lesions and associated disability accumulation largely unaffected [43]. In a 10-year follow-up of patients treated with ocrelizumab, an anti-CD20 B-cell depleting agent, it was shown that PIRA is the leading cause of disability rather than overt relapses [44]. These observations indicate that immunopathology and neurodegeneration act as parallel, mutually reinforcing processes rather than a simple linear cascade. The “inside-out” hypothesis reinforces this view by positing that primary oligodendrocyte or neuronal loss exposes myelin antigens that subsequently stimulate peripheral autoimmunity [45]. Taken together, the evidence argues that durable disease interception will require multipronged therapies capable of modulating immune and neurodegenerative pathways simultaneously. To nominate such combinable targets, we applied an integrated transcriptomic network workflow to MS PBMCs and MS brain-lesion tissue, to identify druggable hubs that can be targeted in concert.

In our research, we used microarray data to characterize DEGs in PBMCs and brain lesion tissue from MS patients and controls. Following uniform pre-processing and stringent filters (adjusted P < 0.05; |log□ fold-change| ≥ 1.5), 142 DEGs were characterized in PBMCs and 1493 in brain. CytoHubba ranking identified a peripherally B-cell–receptor axis (BLK, CD22, CD79A, FCRL1, FCRL5, FCRLA, MS4A1, NIBAN3, TCL1A, and VPREB3) and a glutamatergic-synaptic axis centrally (DLGAP1, DLGAP2, GRIA1, GRIA2, GRIN1, GRIN2A, HOMER1, NLGN1, NRXN1, and SHANK2). Importantly, we identified several modules per compartment, revealing the functions of B-cell signalling and erythrocyte metabolism in blood and excitatory synapse and axon ensheathment in brain. Finally, we identified icotinib and niraparib as high-affinity ligands for the two most important hub genes, CD79A and GRIA2, and thus nominating tractable leads for combination therapy that can target peripheral immune activation and central excitotoxicity in concert.

The first cluster obtained from PBMCs network was mainly related to B cell activation, signaling, proliferation, and regulation. The genes associated with this cluster were upregulated in the PBMCs of MS patients relative to controls, indicating that B cell activation is one of the most important events in the immunopathogenesis of MS. This observation was further supported by the significant upregulation of the purple module in WGCNA of PBMCs transcriptome, which included genes that encode B-cell receptor signaling components (BLNK, CD22, NIBAN3, BANK1, FCRL1, and FCRL2). In support of this, an increased immunoglobulin synthesis rate and abnormal presence of oligoclonal bands patterns of immunoglobulin G and immunoglobulin M in the cerebrospinal fluid (CSF) of MS patients, indicates that B cells are actively involved in MS [46]. Apart from their secreted antibody, soluble products isolated from B cells of MS patients have been shown to contain factors that are toxic to the oligodendrocytes [46]. B cells may mainly contribute to MS through antigen presentation, activation of T cells, and secretion of pro-inflammatory cytokines. Importantly, B cell antigen presentation is crucial in MS as mice lacking the B cell-specific major histocompatibility complex II do not recapitulate anti-myelin oligodendrocyte glycoprotein evoked experimental autoimmune encephalopathy (EAE) [46]. In addition, B cells secrete IL6, which activates Th17 cells that are known as the main pathogenic T cells involved in MS.

In the second cluster we have identified genes that are related to oxygen and carbon dioxide transport and hemoglobin metabolism. The genes related to this were upregulated in MS patients, indicating that erythrocytes and their function are involved in MS. This is supported by observations that erythrocytes in MS patients have impaired antioxidant capacity as evidenced by disturbances in antioxidant enzymes and alterations in blood fluidity [47]. In addition, erythrocytes from MS patients are hypermetabolic and produce high adenosine triphosphate (ATP) levels leading to increased ATP release that induces endothelial cells nitric oxide (NO) production, which disrupts the BBB and causes neuronal injury [48]. The increased NO level interferes with oligodendrocytes DNA and directly causes demyelination and axonal degeneration, thus actively contributing to MS progression [48].

In the third cluster we mainly identified genes associated with the spliceosome. Interestingly, all third cluster genes were upregulated in MS patients compared to healthy controls, except ALYREF, which was downregulated in MS patients. ALYREF encodes Aly/REF export factor, a nuclear protein that functions as a chaperone during mRNA splicing, suggesting that ALYREF downregulation might be interfering with the function of the spliceosome. Increased alternative splicing was noted in MS patients leading to increased immune system reaction to self-antigens [49]. In support of this, several gene polymorphisms associated with alternative splicing of pre-mRNA were found to be increased in B cells of MS patients compared to controls [49–51]. One of the most established consequences of alternative splicing in MS is the skipping of exon 6, which encodes the transmembrane domain of the interleukin 7 receptor (IL7R), leading to an increased secretion of IL7R soluble form [49, 51]. Membrane bound IL7R signaling is required to support homeostatic proliferation of T cells in the thymus and periphery; therefore, excessive soluble IL7R deprives T cells from membrane bound IL7R signaling and this can lead to increased autoreactive clones and also interfere with T cell function, rendering T cells unable to clear viral infections such as the Epstein Barr virus, which is known to be associated with MS [49]. The importance of the spliceosome was further corroborated by the upregulation of the blue WGCNA module, which contained genes associated with mRNA splicing.

The final PBMCs cluster was associated with immune cell activation and response to molecules of bacterial origin, which is crucial in the genesis of MS. Among the 12 genes associated with the fourth cluster, three genes (STAT2, CLEC7A, and ANXA1) were consistently downregulated in MS patients compared to controls. STAT2 is a transcriptional factor known to be involved in type I/III interferon signaling, and its downregulation in MS patients may indicate reduced interferon signaling leading to overactivation of the immune system, increased inflammation, disruption of BBB, and neuronal injury [52]. CLEC7A encodes dectin-1, a c-type lectin-like receptor, which recognizes β-glucan carbohydrates present in fungi. In this study, CLEC7A was consistently downregulated in MS patients’ PBMCs and this was in line with a previous finding that demonstrated that mice deficient in dectin-1 develop severe MS symptoms upon induction of EAE [53]. This heightened EAE symptoms compared to wild-type mice is mainly due Th17 cells activation and reduced T cell regulatory cells development, suggesting that dectin-1 may be a target that upon its activation may reduce autoimmune damage in MS patients. The last downregulated gene in MS patients is ANXA1, which encodes annexin A1, a glucocorticoid-responsive protein that reduces inflammation. Reduced ANXA1 secretion in MS patients is associated with increased T cell activation, proliferation, and metabolism and correlated with increased disease severity and inflammation [54]. The dysregulation of the immune system is further supported by the downregulation of the brown WGCNA module, which signifies reduced natural killer cells activity, that has important immunoregulatory function.

The first MS brain lesion cluster was associated with glutamatergic signaling and it included genes such as (GRIA1, GRIA2, GRIN1, and GRIN2A), which encode different ionotropic glutamate receptors. These genes were upregulated in MS brain lesions, mirroring classic histopathological and *in vivo* findings implicating glutamate excitotoxicity in MS oligodendrocyte and axonal damage [55]. Immunostaining examination of MS post-mortem patients brain samples has revealed an increased expression of the calcium permeable GluA1 subunit of AMPA receptor on oligodendrocytes [56]. Interestingly the first cluster included an increased mitogen-activated protein kinase signal transduction that may play a role in the upregulation of glutamate receptors on oligodendrocytes. In addition, glutamate levels are elevated in CSF of MS patients and its elevation correlated positively with disease activity and extension of axonal damage [55]. The role of increased glutamatergic signaling is supported by our gene enrichment results pinpointing positive regulation of excitatory postsynaptic membrane potential at the glutamatergic synapse. Moreover, the GSEA result showed an upregulation of synaptic signaling, increased transmission across chemical synapses, and excessive neuroinflammation and glutamatergic signaling. Together, these data suggest that an excessive increase in glutamatergic signaling can lead to neuronal excitotoxicity, oligodendrocytes loss, and demyelination in MS.

The second cluster was related to neurons and axon ensheathment and these processes were downregulated according to our GSEA results. This cluster also identified voltage-gated sodium channels as important compartments and voltage-gated sodium channel activity as an important function. This is in line with findings showing changed expression of sodium channel Na_v_1.6 along extensive regions of demyelinated axons in MS plaques [57]. The persistent sodium current by Na_v_1.6 drives colocalized Na^+^/Ca^2+^ exchanger to import excessive Ca^2+^ levels into axons that causes mitochondrial dysfunction leading to axonal damage and degeneration associated with MS. The GSEA results also pointed to the downregulation of lipids and steroids metabolism and reduced oligodendrocytes differentiation, which are critical for remyelination. These findings suggest ionic and metabolic bottlenecks converge to interfere with axonal ensheathment and oligodendrocyte differentiation in MS.

The third cluster is mainly related to netrin-activated pathway signaling, which is involved in neuronal migration and axon guidance. The dysregulation of the netrin-activated pathway in MS interferes with its BBB stabilization activity and skews microglia towards a pro-inflammatory phenotype, leading to an increase in neuroinflammation. Netrin-1 injection in the EAE model of MS has been shown to reduce BBB disruption and decrease both clinical and pathological disease severity indices [58]. The final brain cluster was enriched in protein import into nucleus, nuclear-pore complex, and nuclear localization-sequence binding. Direct MS-specific evidence is lacking, but nucleocytoplasmic transport deficits are a proven early etiology of neuronal vulnerability in other neurodegenerative diseases [59, 60]. The dysregulation of nucleocytoplasmic transport associated with MS lesions suggests that transport defects may place neurons in an irreversible stress state.

LASSO identified three PBMCs hub genes that were all related to B cell cluster, including (FCRL1, CD22, and CD79A). CD79A has been recently shown to be able to predict the risk of MS and was also upregulated in an EAE model of MS [61]. Moreover, Mendelian randomization using the inverse variance weighted fixed effects estimate confirmed the increased CD79A expression is linked to an increased risk of MS development, suggesting that CD79A can be used as a diagnostic and therapeutic target for MS [61]. Therefore, targeting CD79A may prove to be an effective way to reduce B cell activation in MS. CD79A is upstream of BTK and BTK inhibitors are an emerging class of therapeutics for MS and numerous preclinical studies have shown that its inhibition suppresses B cell activation and reduces the pathological features of MS [62, 63]. Moreover, CD79A-targeted therapies can induce transient anergy in autoreactive B cells by uncoupling the BCR-induced tyrosine phosphorylation and calcium mobilization without causing adverse effects associated with B cell depleting therapies [64]. The effectiveness of CD79A targeting is further reinforced by preventing autoimmunity in EAE without causing significant B cell depletion, confirming the importance of CD79A targeting in MS [64]. The top three CytoHubba brain lesion genes were GRIN2A, NRXN1, and GRIA2. GRIA2 encodes the GluA2 subunit of AMPA receptors, and selective modulation of GluA2 with ZCAN262 has been shown to significantly attenuate disease severity in both the EAE and cuprizone□induced demyelination models of MS [65]. Other studies have also shown that modulation of AMPA glutamate receptor reduces demyelination in EAE mouse model of MS [66–68]. Docking and 200 ns molecular dynamics simulations showed that both icotinib and niraparib have high-affinity binding to CD79A and GRIA2. Dual-target engagement of these two drugs at two targets suggests them as repurposing candidates capable of modulating B cell receptor signaling and excitotoxic glutamatergic pathways concurrently in MS.

Despite the integrative pipeline employed, several limitations merit consideration. First, the reliance on microarray datasets of modest size (14 MS and 15 control PBMC samples; 5 MS and 2 control brain lesions) may introduce sampling bias and platform-specific artefacts; replication of key signatures in larger, RNA sequencing–based cohorts is essential. Second, network and machine-learning approaches infer statistical associations rather than causality; hub-gene expression and module membership should be confirmed by quantitative PCR or immunohistochemistry. Third, several highly upregulated transcripts in the PBMC dataset, such as hemoglobin subunits, likely reflect residual red blood cell or reticulocyte contamination during PBMCs preparation. This recognized artifact in blood transcriptomics should be considered when interpreting these results. Fourth, the *in silico* identification of icotinib and niraparib as dual CD79A/GRIA2 binders offers a promising repurposing hypothesis, but requires validation through dose-response studies in established MS models. Future investigations should (i) validate CD79A, GRIA2 at the protein level in independent human samples; (ii) employ genetic perturbation (e.g., CRISPR/Cas9) in primary B cells and oligodendrocyte precursor cultures to elucidate mechanistic contributions to MS pathology; and (iii) perform *in vivo* evaluation of icotinib and niraparib in EAE and cuprizone models to determine efficacy, optimal dosing and safety. Addressing these tasks will be critical for advancing the proposed dual-target repurposing strategy toward clinical translation.

## 5. Conclusion

This study demonstrates that an integrated transcriptomic network approach can resolve the complex, multi-axis pathology of multiple sclerosis by uncovering hubs in both peripheral blood and CNS lesions. In PBMCs, we identified a B cell receptor module anchored on CD79A, alongside erythrocyte metabolic and spliceosome programmes, while lesion transcriptomes revealed a glutamatergic synapse hub centered on GRIA2 and modules implicating axonal ensheathment, netrin-1 signalling and nucleocytoplasmic transport. Machine-learning prioritization confirmed FCRL1, CD22, and CD79A as important blood markers. Critically, virtual screening and 200 ns molecular dynamics simulations showed that icotinib and niraparib can engage both CD79A and GRIA2, providing a repurposing strategy to co-modulate peripheral autoimmunity and central excitotoxicity. To translate this, future work must validate hub expression in independent cohorts, perform mechanistic perturbations in primary cells, and assess efficacy and safety of the proposed drug combinations in EAE and cuprizone models. By moving beyond single-mechanism paradigms, this dual-compartment, repurposing-focused framework holds promise to overcome the limitations of current monotherapies and achieve durable disease interception in MS.

## References

1. Compston A, Winedl H, Kieseier B. Coles. Multiple sclerosis Lancet. 2008;372:1502–17.

2. Trapp BD, Nave K-A. Multiple sclerosis: an immune or neurodegenerative disorder? Annu Rev Neurosci. 2008;31(1):247–69.

3. Hauser SL, Oksenberg JR. The neurobiology of multiple sclerosis: genes, inflammation, and neurodegeneration. Neuron. 2006;52(1):61–76.

4. Barnett MH, Prineas JW. Relapsing and remitting multiple sclerosis: pathology of the newly forming lesion. Annals of neurology. 2004;55(4):458–68.

5. Lucchinetti C, Brück W, Parisi J, Scheithauer B, Rodriguez M, Lassmann H. Heterogeneity of multiple sclerosis lesions: implications for the pathogenesis of demyelination. Annals of Neurology: Official Journal of the American Neurological Association and the Child Neurology Society. 2000;47(6):707–17.

6. Hauser SL, Waubant E, Arnold DL, Vollmer T, Antel J, Fox RJ, et al. B-cell depletion with rituximab in relapsing–remitting multiple sclerosis. New England Journal of Medicine. 2008;358(7):676–88.

7. Bellanca CM, Augello E, Mariottini A, Bonaventura G, La Cognata V, Di Benedetto G, et al. Disease Modifying Strategies in Multiple Sclerosis: New Rays of Hope to Combat Disability? Current Neuropharmacology. 2024;22(8):1286–326.

8. Langfelder P, Horvath S. WGCNA: an R package for weighted correlation network analysis. BMC bioinformatics. 2008;9(1):559.

9. Szklarczyk D, Gable AL, Lyon D, Junge A, Wyder S, Huerta-Cepas J, et al. STRING v11: protein–protein association networks with increased coverage, supporting functional discovery in genome-wide experimental datasets. Nucleic acids research. 2019;47(D1):D607–D13.

10. Pushpakom S, Iorio F, Eyers PA, Escott KJ, Hopper S, Wells A, et al. Drug repurposing: progress, challenges and recommendations. Nature reviews Drug discovery. 2019;18(1):41–58.

11. Davis S, Meltzer PS. GEOquery: a bridge between the Gene Expression Omnibus (GEO) and BioConductor. Bioinformatics. 2007;23(14):1846–7.

12. Huber W, Carey VJ, Gentleman R, Anders S, Carlson M, Carvalho BS, et al. Orchestrating high-throughput genomic analysis with Bioconductor. Nature methods. 2015;12(2):115–21.

13. Ritchie ME, Phipson B, Wu D, Hu Y, Law CW, Shi W, et al. limma powers differential expression analyses for RNA-sequencing and microarray studies. Nucleic acids research. 2015;43(7):e47-e.

14. Leek JT, Storey JD. Capturing heterogeneity in gene expression studies by surrogate variable analysis. PLoS genetics. 2007;3(9):e161.

15. Ritchie ME, Silver J, Oshlack A, Holmes M, Diyagama D, Holloway A, et al. A comparison of background correction methods for two-colour microarrays. Bioinformatics. 2007;23(20):2700–7.

16. Wilkinson L. ggplot2: elegant graphics for data analysis by Wickham, H. Oxford University Press; 2011.

17. Szklarczyk D, Kirsch R, Koutrouli M, Nastou K, Mehryary F, Hachilif R, et al. The STRING database in 2023: protein–protein association networks and functional enrichment analyses for any sequenced genome of interest. Nucleic acids research. 2023;51(D1):D638–D46.

18. Shannon P, Markiel A, Ozier O, Baliga NS, Wang JT, Ramage D, et al. Cytoscape: a software environment for integrated models of biomolecular interaction networks. Genome research. 2003;13(11):2498–504.

19. Men X, Shi X, Xu Q, Liu M, Yang H, Wang L, et al. Exploring the pathogenesis of chronic atrophic gastritis with atherosclerosis via microarray data analysis. Medicine. 2024;103(16):e37798.

20. Csardi G, Nepusz T. The igraph software. Complex syst. 2006;1695:1–9.

21. Pedersen T. An implementation of grammar of graphics for graphs and networks [R Package Ggraph Version 2.0. 4]. Corpus ID. 2020;22910764.

22. Yu G, Wang L-G, Han Y, He Q-Y. clusterProfiler: an R package for comparing biological themes among gene clusters. Omics: a journal of integrative biology. 2012;16(5):284–7.

23. Yu G, He Q-Y. ReactomePA: an R/Bioconductor package for reactome pathway analysis and visualization. Molecular BioSystems. 2016;12(2):477–9.

24. Dolgalev I. msigdbr: MSigDB gene sets for multiple organisms in a tidy data format. R package version. 2022;7(1).

25. Zhang B, Horvath S. A general framework for weighted gene co-expression network analysis. Statistical applications in genetics and molecular biology. 2005;4(1).

26. Langfelder P, Zhang B, Horvath S. Defining clusters from a hierarchical cluster tree: the Dynamic Tree Cut package for R. Bioinformatics. 2008;24(5):719–20.

27. Tibshirani R. Regression shrinkage and selection via the lasso. Journal of the Royal Statistical Society Series B: Statistical Methodology. 1996;58(1):267–88.

28. Friedman JH, Hastie T, Tibshirani R. Regularization paths for generalized linear models via coordinate descent. Journal of statistical software. 2010;33(1):1–22.

29. Kuhn M. Building predictive models in R using the caret package. Journal of statistical software. 2008;28:1–26.

30. Robin X, Turck N, Hainard A, Tiberti N, Lisacek F, Sanchez J-C, et al. pROC: an open-source package for R and S+ to analyze and compare ROC curves. BMC bioinformatics. 2011;12(1):77.

31. Pettersen EF, Goddard TD, Huang CC, Couch GS, Greenblatt DM, Meng EC, et al. UCSF Chimera—a visualization system for exploratory research and analysis. Journal of computational chemistry. 2004;25(13):1605–12.

32. Lipinski CA. Lead-and drug-like compounds: the rule-of-five revolution. Drug discovery today: Technologies. 2004;1(4):337–41.

33. Veber DF, Johnson SR, Cheng H-Y, Smith BR, Ward KW, Kopple KD. Molecular properties that influence the oral bioavailability of drug candidates. Journal of medicinal chemistry. 2002;45(12):2615–23.

34. Daina A, Michielin O, Zoete V. SwissADME: a free web tool to evaluate pharmacokinetics, drug-likeness and medicinal chemistry friendliness of small molecules. Scientific reports. 2017;7(1):42717.

35. Tian H, Ketkar R, Tao P. ADMETboost: a web server for accurate ADMET prediction. Journal of molecular modeling. 2022;28(12):408.

36. Tian H, Xiao S, Jiang X, Tao P. PASSer: fast and accurate prediction of protein allosteric sites. Nucleic Acids Research. 2023;51(W1):W427–W31.

37. Dallakyan S, Olson AJ. Small-molecule library screening by docking with PyRx. Chemical biology: methods and protocols: Springer; 2014. p. 243–50.

38. Abraham MJ, Murtola T, Schulz R, Páll S, Smith JC, Hess B, et al. GROMACS: High performance molecular simulations through multi-level parallelism from laptops to supercomputers. SoftwareX. 2015;1:19–25.

39. Miller III BR, McGee Jr TD, Swails JM, Homeyer N, Gohlke H, Roitberg AE. MMPBSA. py: an efficient program for end-state free energy calculations. Journal of chemical theory and computation. 2012;8(9):3314–21.

40. Valdés-Tresanco MS, Valdés-Tresanco ME, Valiente PA, Moreno E. gmx_MMPBSA: a new tool to perform end-state free energy calculations with GROMACS. Journal of chemical theory and computation. 2021;17(10):6281–91.

41. Wootla B, Eriguchi M, Rodriguez M. Is multiple sclerosis an autoimmune disease? Autoimmune diseases. 2012;2012(1):969657.

42. Coles AJ, Cohen JA, Fox EJ, Giovannoni G, Hartung H-P, Havrdova E, et al. Alemtuzumab CARE-MS II 5-year follow-up: efficacy and safety findings. Neurology. 2017;89(11):1117–26.

43. Maggi P, Vanden Bulcke C, Pedrini E, Bugli C, Sellimi A, Wynen M, et al. B cell depletion therapy does not resolve chronic active multiple sclerosis lesions. EBioMedicine. 2023;94.

44. Ingwersen J, Masanneck L, Pawlitzki M, Samadzadeh S, Weise M, Aktas O, et al. Real-world evidence of ocrelizumab-treated relapsing multiple sclerosis cohort shows changes in progression independent of relapse activity mirroring phase 3 trials. Scientific Reports. 2023;13(1):15003.

45. Sen MK, Almuslehi MS, Shortland PJ, Coorssen JR, Mahns DA. Revisiting the pathoetiology of multiple sclerosis: has the tail been wagging the mouse? Frontiers in immunology. 2020;11:572186.

46. Li R, Patterson KR, Bar-Or A. Reassessing B cell contributions in multiple sclerosis. Nature immunology. 2018;19(7):696–707.

47. Groen K, Maltby VE, Sanders KA, Scott RJ, Tajouri L, Lechner-Scott J. Erythrocytes in multiple sclerosis–forgotten contributors to the pathophysiology? Multiple Sclerosis Journal– Experimental, Translational and Clinical. 2016;2:2055217316649981.

48. Geiger M, Hayter E, Martin R, Spence D. Red blood cells in type 1 diabetes and multiple sclerosis and technologies to measure their emerging roles. Journal of Translational Autoimmunity. 2022;5:100161.

49. Evsyukova I, Somarelli JA, Gregory SG, Garcia-Blanco MA. Alternative splicing in multiple sclerosis and other autoimmune diseases. RNA biology. 2010;7(4):462–73.

50. Putscher E, Hecker M, Fitzner B, Boxberger N, Schwartz M, Koczan D, et al. Genetic risk variants for multiple sclerosis are linked to differences in alternative pre-mRNA splicing. Frontiers in Immunology. 2022;13:931831.

51. Hecker M, Ruege A, Putscher E, Boxberger N, Rommer PS, Fitzner B, et al. Aberrant expression of alternative splicing variants in multiple sclerosis–A systematic review. Autoimmunity reviews. 2019;18(7):721–32.

52. Manoochehrabadi S, Arsang-Jang S, Mazdeh M, Inoko H, Sayad A, Taheri M. Analysis of STAT1, STAT2 and STAT3 mRNA expression levels in the blood of patients with multiple sclerosis. Human Antibodies. 2019;27(2):91–8.

53. Yan C, Tang N, Guo H, Zhang J. C-Type lectin receptor dectin-1 suppresses the development of experimental autoimmune encephalomyelitis. The Journal of Immunology. 2020;204(1_Supplement):150.19–.19.

54. Colamatteo A, Maggioli E, Azevedo Loiola R, Hamid Sheikh M, Calì G, Bruzzese D, et al. Reduced annexin A1 expression associates with disease severity and inflammation in multiple sclerosis patients. The Journal of Immunology. 2019;203(7):1753–65.

55. Blaylock RL. Immunoexcitoxicity as the possible major pathophysiology behind multiple sclerosis and other autoimmune disorders. Surgical Neurology International. 2025;16:26.

56. Newcombe J, Uddin A, Dove R, Patel B, Turski L, Nishizawa Y, et al. Glutamate receptor expression in multiple sclerosis lesions. Brain pathology. 2008;18(1):52–61.

57. Craner MJ, Newcombe J, Black JA, Hartle C, Cuzner ML, Waxman SG. Molecular changes in neurons in multiple sclerosis: altered axonal expression of Nav1. 2 and Nav1. 6 sodium channels and Na+/Ca2+ exchanger. Proceedings of the National Academy of Sciences. 2004;101(21):8168–73.

58. Podjaski C, Alvarez JI, Bourbonniere L, Larouche S, Terouz S, Bin JM, et al. Netrin 1 regulates blood–brain barrier function and neuroinflammation. Brain. 2015;138(6):1598–612.

59. Spead O, Zaepfel BL, Rothstein JD. Nuclear pore dysfunction in neurodegeneration. Neurotherapeutics. 2022;19(4):1050–60.

60. Gleixner AM, Verdone BM, Otte CG, Anderson EN, Ramesh N, Shapiro OR, et al. NUP62 localizes to ALS/FTLD pathological assemblies and contributes to TDP-43 insolubility. Nature communications. 2022;13(1):3380.

61. Ding S, Zhang Y, Tang Y, Zhang Y, Liu M. Combining gene expression microarrays and Mendelian randomization: exploring key immune-related genes in multiple sclerosis. Frontiers in Neurology. 2024;15:1437778.

62. Montalban X, Arnold DL, Weber MS, Staikov I, Piasecka-Stryczynska K, Willmer J, et al. Placebo-controlled trial of an oral BTK inhibitor in multiple sclerosis. New England Journal of Medicine. 2019;380(25):2406–17.

63. Correale J. BTK inhibitors as potential therapies for multiple sclerosis. The Lancet Neurology. 2021;20(9):689–91.

64. Wemlinger SM, Parker Harp CR, Yu B, Hardy IR, Seefeldt M, Matsuda J, et al. Preclinical analysis of candidate anti-human CD79 therapeutic antibodies using a humanized CD79 mouse model. The Journal of Immunology. 2022;208(7):1566–84.

65. Zhai D, Yan S, Samsom J, Wang L, Su P, Jiang A, et al. Small-molecule targeting AMPA-mediated excitotoxicity has therapeutic effects in mouse models for multiple sclerosis. Science Advances. 2023;9(49):eadj6187.

66. Mey GM, Evonuk KS, Shelestak J, Irfan M, Wolfe LM, Laye SE, et al. Inhibiting AMPA receptor signaling in oligodendrocytes rescues synapse loss in a model of autoimmune demyelination. iScience. 2024;27(11).

67. Sindi M, Dietrich M, Klees D, Gruchot J, Hecker C, Silbereis J, et al. Positive allosteric modulation of AMPA receptors via PF4778574 leads to reduced demyelination and clinical disability in experimental models of multiple sclerosis. Frontiers in Immunology. 2025;16:1532877.

68. Smith T, Groom A, Zhu B, Turski L. Autoimmune encephalomyelitis ameliorated by AMPA antagonists. Nature medicine. 2000;6(1):62–6.

