## Supplementary material for "Integrated Blood–Brain Transcriptomics Reveals CD79A and GRIA2 as Drug Repurposing Targets in Multiple Sclerosis": Supplementary Material.docx

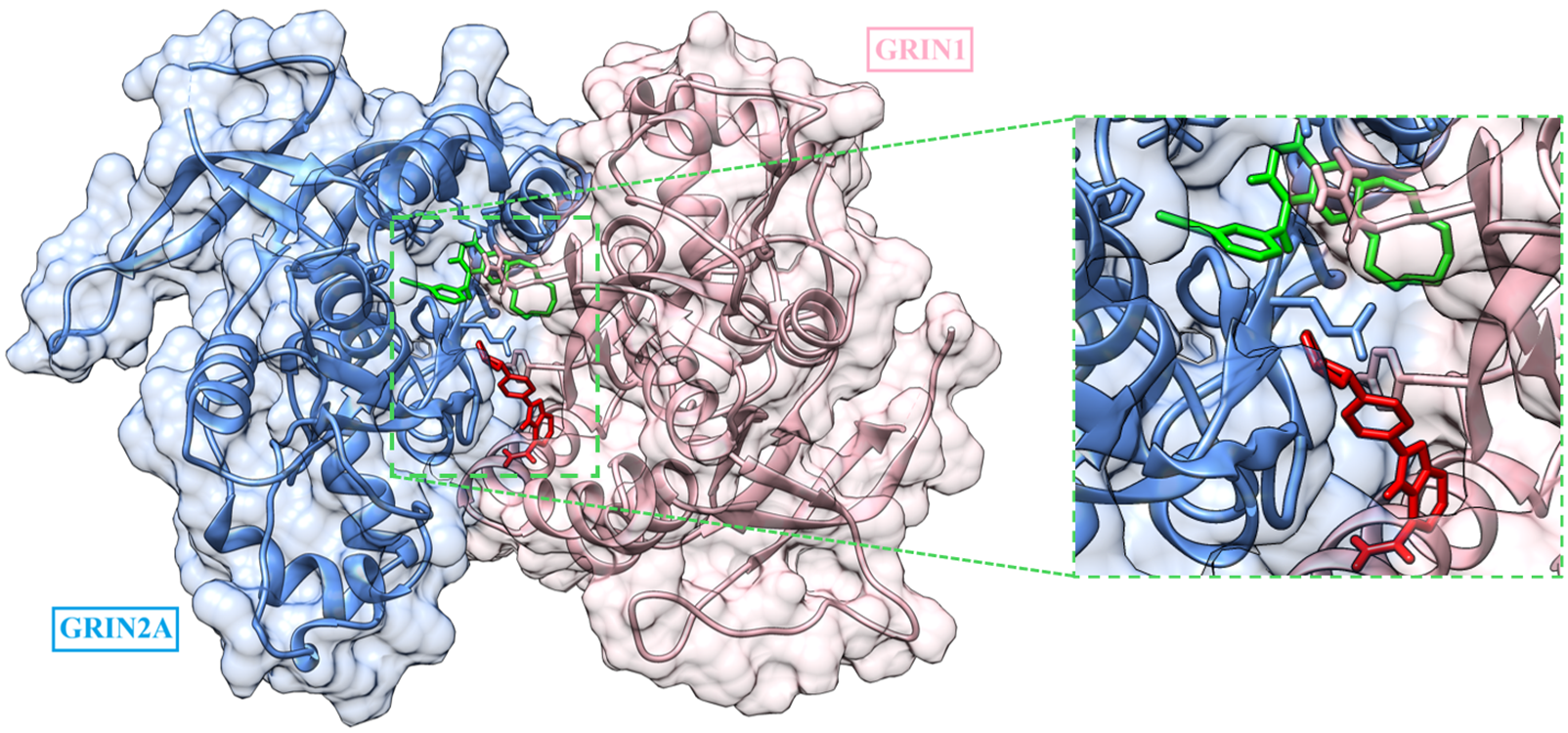

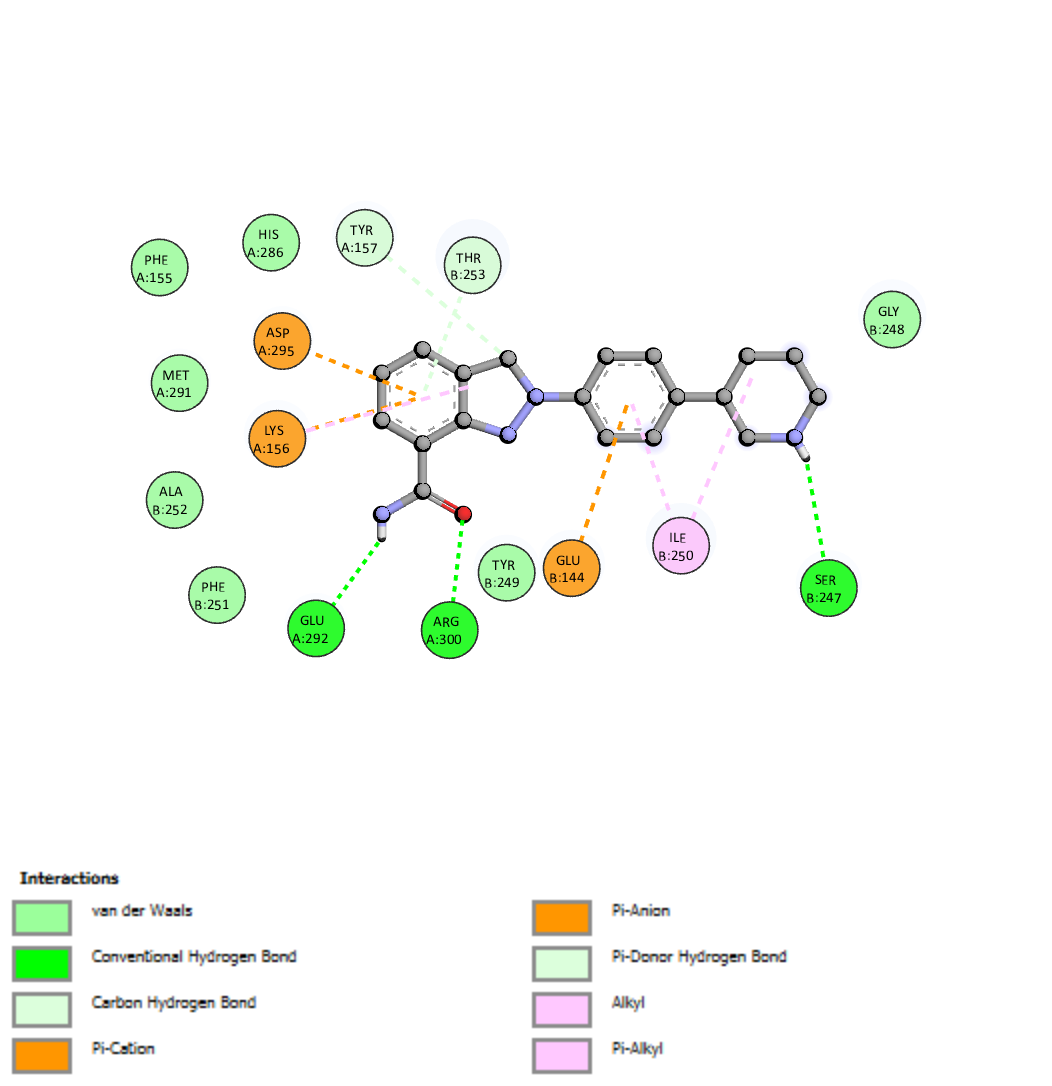

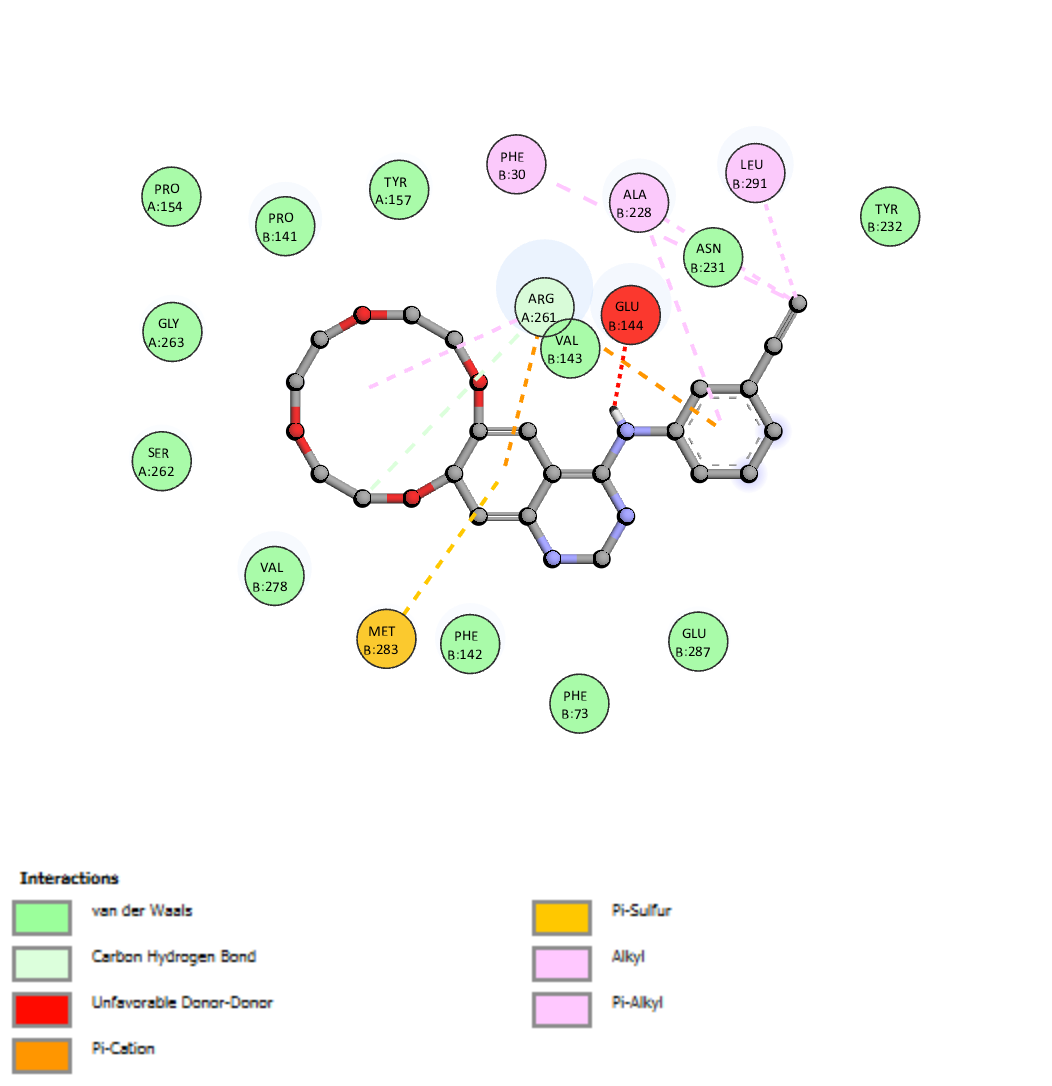

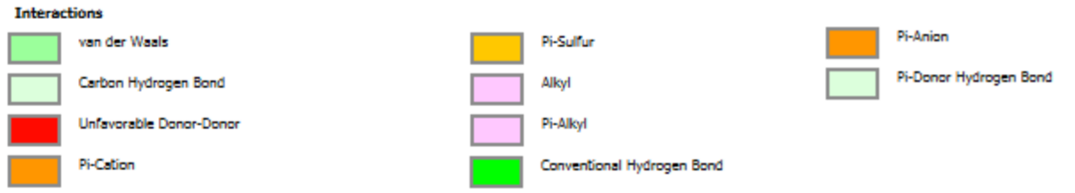

**[C]**

**[B]**

**[A]**

**Figure S1.** Molecular docking of compounds against GluN1/GluN2A. **(A)** The 3D binding configuration of Icotinib (Green) and Niraparib (Red). The molecular interaction fingerprints in 2D showed that atoms of Icotinib **(B)** and Niraparib **(C)** interacted with amino acid residues important for binding with GluN1/GluN2A.

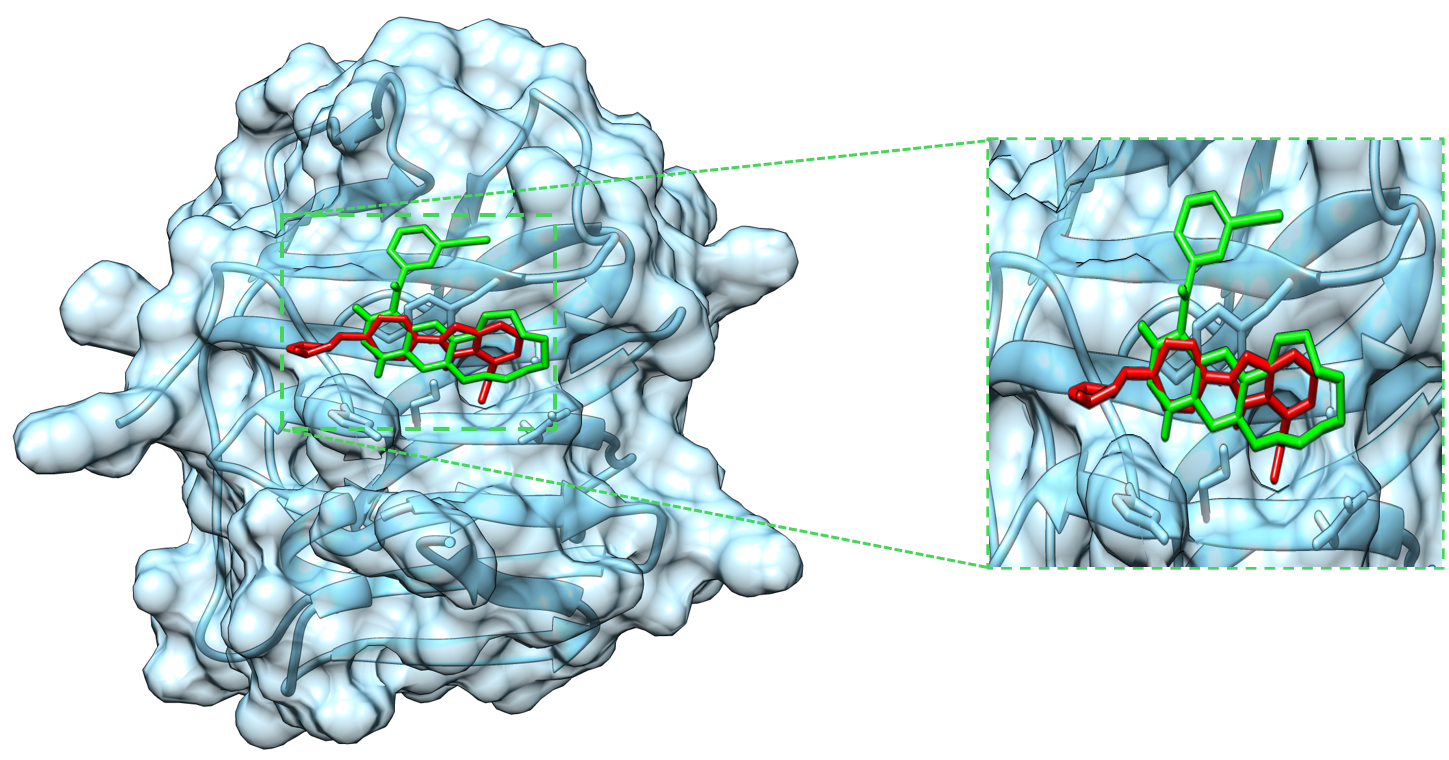

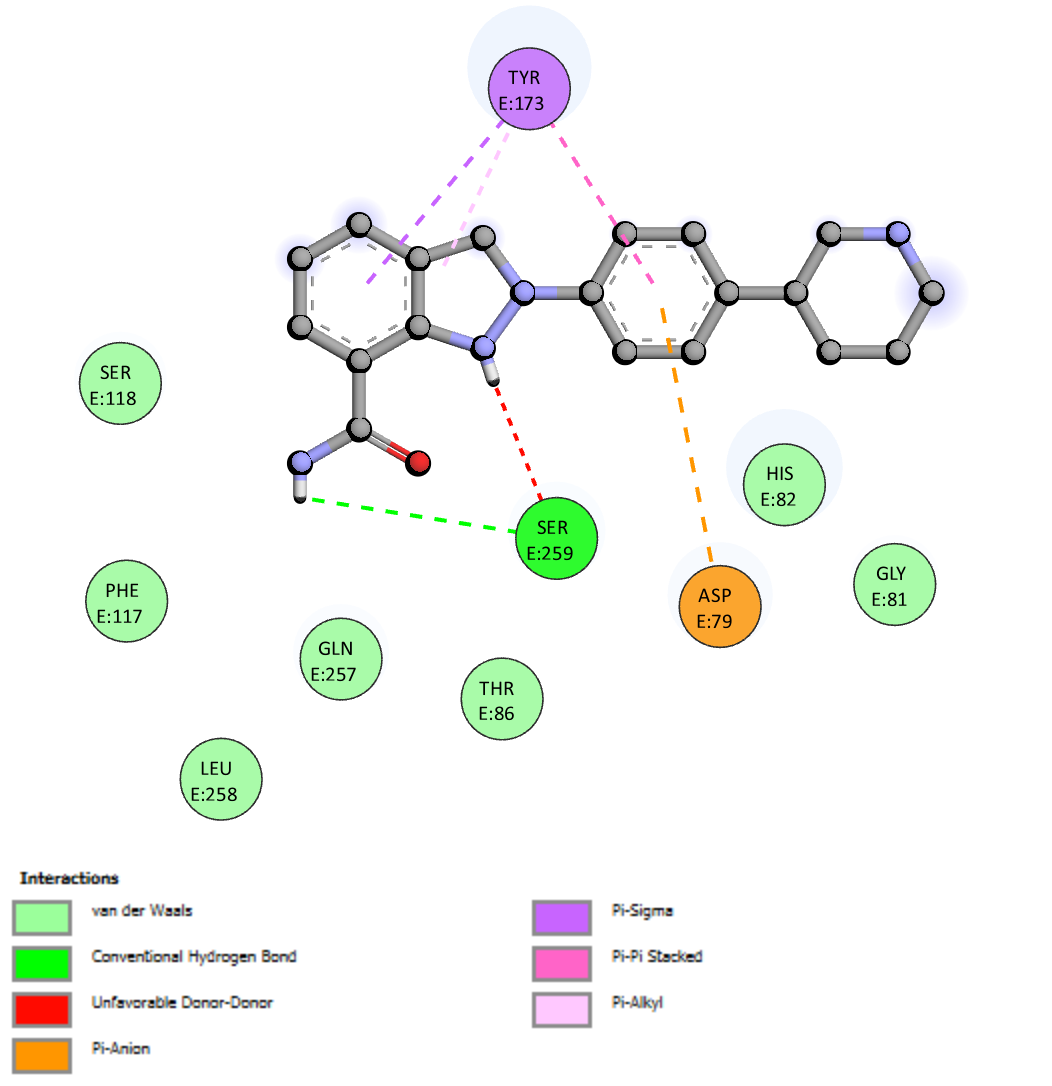

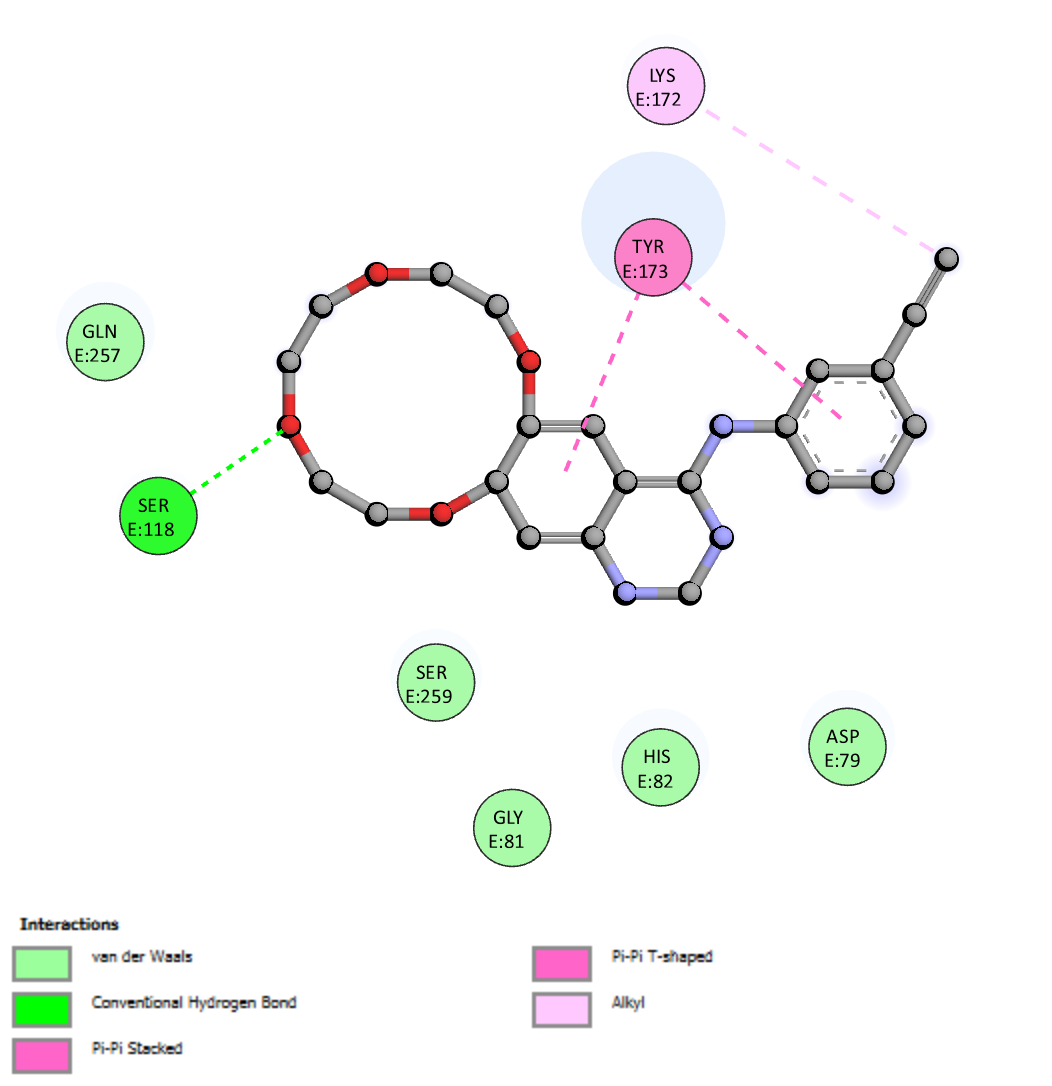

**[C]**

**[B]**

**[A]**

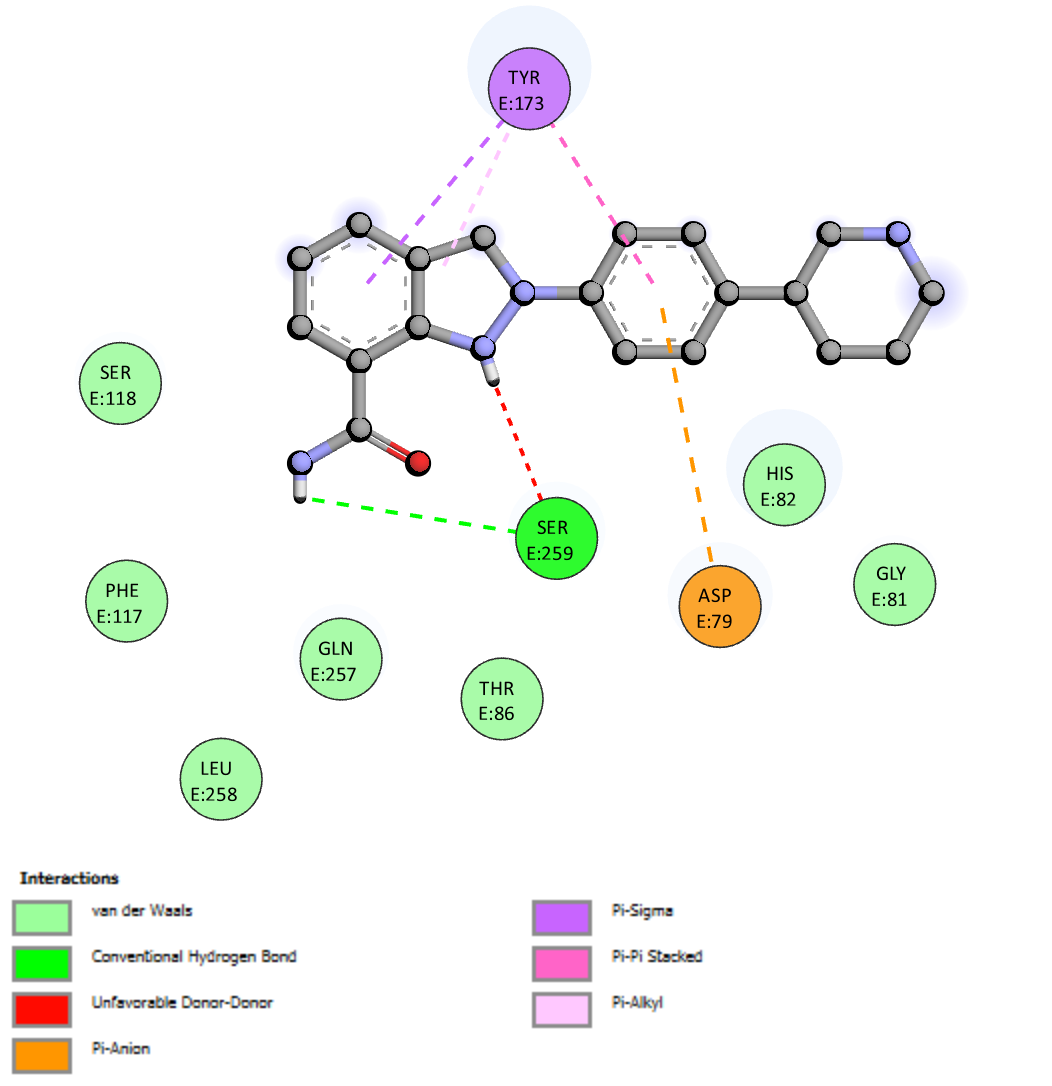

**Figure S2.** Molecular docking of compounds against NRXN1. **(A)** The 3D binding configuration of Icotinib (Green) and Niraparib (Red). The molecular interaction fingerprints in 2D showed that atoms of Icotinib **(B)** and Niraparib **(C)** interacted with amino acid residues important for binding with NRXN1.

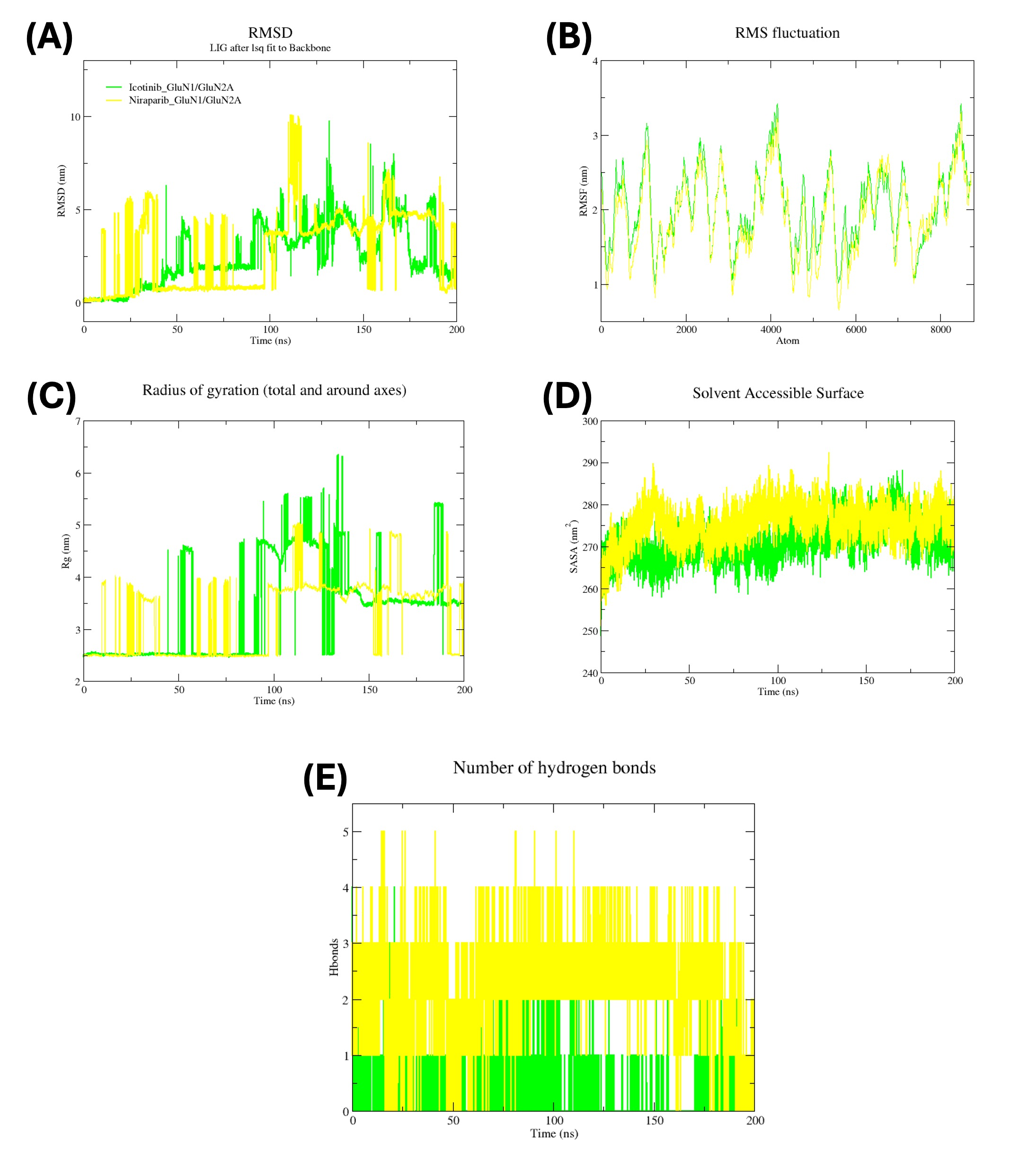

**Figure S3. Molecular dynamics simulation analysis of Icotinib and Niraparib complexes with GRIN1/GRIN2A:** **(A)** RMSD, **(B)** RMSF, **(C)** Radius of Gyration, **(D)** Solvent Accessible Surface Area, and **(E)** Hydrogen Bond Profiles Over 200 ns Simulations.

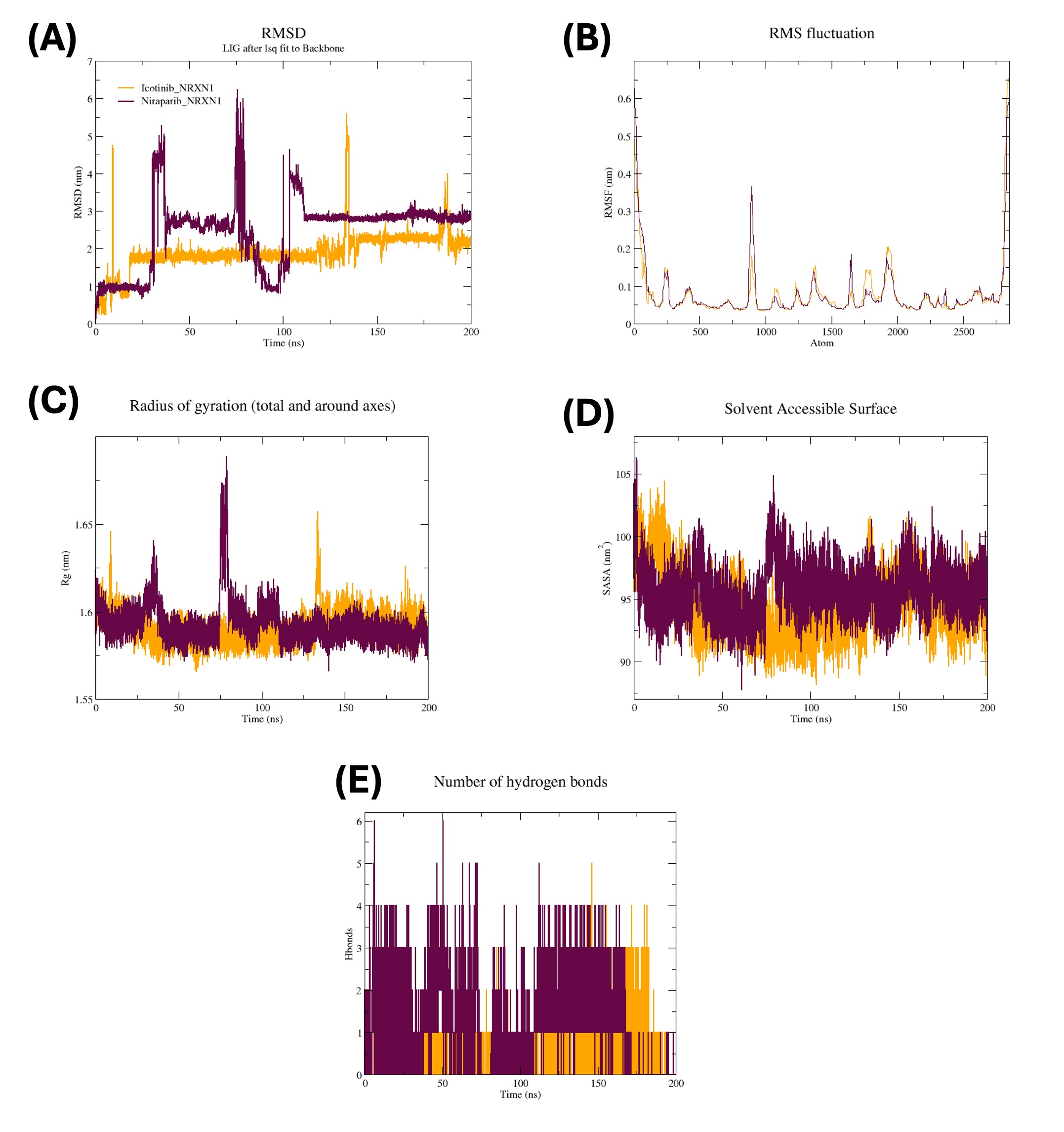

**Figure S4.** **Molecular dynamics simulation analysis of Icotinib and Niraparib complexes with NRXN1:** **(A)** RMSD, **(B)** RMSF, **(C)** Radius of Gyration, **(D)** Solvent Accessible Surface Area, and **(E)** Hydrogen Bond Profiles Over 200 ns Simulations.
